# The 3D Genome of *Gigaspora margarita* Unveils Stable Chromatin and Nucleolar Organization and Symbiont-Dependent Genome Dynamics

**DOI:** 10.64898/2026.01.14.699541

**Authors:** Ken Mugambi, Jordana Oliveira, Franco Magurno, Alessandra Salvioli di Fossalunga, Mara Novero, Luisa Lanfranco, Stefano Ghignone, Gökalp Yildirir, Yan Wang, Paola Bonfante, Nicolas Corradi

## Abstract

Arbuscular mycorrhizal fungi (AMF) are widespread plant symbionts that enhance nutrient acquisition and influence ecosystem productivity. Previous chromosome-level assemblies of a model species revealed a two-compartment genome architecture (active A and repressed B chromatin compartments), yet its conservation across evolutionarily distant AMF lineages remains unresolved. Here, we present a chromosome-scale and 3D genome assembly of *Gigaspora margarita* isolate BEG34—the largest and most repeat-rich AMF genome to date—alongside that of its obligate endobacterium, *Candidatus* Glomerobacter gigasporarum (*Ca*Gg), using PacBio HiFi and Hi-C sequencing. The *G. margarita* genome comprises 43 chromosomes (792 Mb) organized into stable A/B compartments and Topologically Associating Domains structures, irrespective of the presence of endobacteria. We uncover 21 divergent rDNA operons distributed across six chromosomes and show that these physically interact, suggesting conserved nucleolar organization. We also reveal that the *Ca*Gg genome is tripartite and mobilome-rich, encoding prophages, an orphan CRISPR array, and complete pathways for many novel and essential cofactors, including heme, which may enhance host bioenergetics. We also find that the endobacterium’s presence regulates transposable elements in *G. margarita*. These findings reveal conserved principles of chromatin architecture in AMF symbionts and highlight the tight molecular interplay between fungal hosts and their endosymbionts, offering new insights into genome evolution and symbiotic adaptation.

## Introduction

Arbuscular mycorrhizal fungi (AMF) are plant root symbionts that belong to the subphylum Glomeromycotina ^1^. As obligate biotrophs, AMF require a living plant host to complete their life cycle, leading them to colonize the cortical root cells and develop specialized tree-like structures called arbuscules ^2^. Within these structures, the plant supplies the fungi with lipids and sugars ^3,4^, while the AMF provides the plant with essential nutrients that are limiting factors for plant growth ^5^, primarily phosphate, resulting in improved crop yields ^6,7^, carbon storage ^8^, and enhanced defence against pathogens ^9^. AMF are always multinucleated, with individual spores carrying up to 20,000 haploid nuclei in some species ^10^. Although sexual reproductive structures have not yet been observed in AMF, genome and single-nucleus analyses have shown that these organisms carry conserved mating-related genes ^11–16^ and follow homokaryotic/heterokaryotic cycles ^17,18^, which define sexual processes in fungi ^19,20^. Genome analyses show that AMF plant dependence is likely linked to the loss of genes involved in fatty acid production, thiamine biosynthesis, sugar utilization, and plant cell wall degradation^12,21^. Their genomes are also highly enriched in transposable elements (TEs), and closely related AMF strains exhibit striking variation in gene content ^22,23^. The abundance of TEs in AMF is the main reason their genome assemblies have long been highly fragmented, thereby hampering our understanding of their genetic structure and overall genome biology.

This issue was recently addressed by combining long reads with chromatin capture (Hi-C) datasets, which allowed the generation of chromosomal-level assemblies for model AMF *R. irregularis* strains (order Glomerales)^18,24^. This approach revealed that AMF chromatin separates into two compartments (A/B). The compartment A contains transcriptionally active genes, high methylation of transposable elements (TEs), and most conserved core genes. In contrast, compartment B harbours transcriptionally repressed genes and is rich in genes encoding secreted proteins, candidate effectors, and TEs that are upregulated *in planta* (vs. extra-radical mycelium) ^18,21,24–28^. This stage-specific upregulation suggests that root colonization leads to the relaxation of the B sub-compartments involved in the molecular dialogues between partners of the mycorrhizal symbiosis ^21,24^. In support of this, transmission electron microscopy of the AMF *Gigaspora margarita* shows shifts in chromatin condensation, transitioning from a tightly packed state in spores to a looser state in intra-radical hyphae during plant root colonization ^29^.

In addition to being separated into A/B compartments, the available *R. irregularis* genomes are also organized into finer-scale structural units known as Topologically Associating Domains (TADs)-like structures, within which chromatin interactions occur more frequently than with neighbouring regions ^24,30–32^. In model eukaryotes, TADs are separated by “boundaries” that act as insulators, limiting epigenetic interactions between adjacent TADs ^30,31,33^, and are often enriched for protein-coding genes and depleted of DNA repeats ^31,33,34^. In contrast, in fungal species, repeats were shown to dominate TAD boundaries^35^, highlighting their functional divergence and potentially higher malleability. Hi-C-based identification of A/B compartments in AMF remains confined to the model AMF species *R. irregularis*, and independent support for their existence and conservation in other AMF lineages is lacking. Similarly, although data from model eukaryotes ^36–38^ supports the hypothesis that the AMF A/B sub-compartments may change conformation in response to host and environmental factors, direct evidence based on chromatin analyses that such changes occur in AMF has not yet been provided.

Here, we aim to fill these knowledge gaps by sequencing the genome of *G. margarita* isolate BEG34 (order Diversisporales) using PacBio HiFi and Hi-C data. This species contains the largest and most repeat-rich AMF genome known to date^39^. It also carries beneficial obligate *Burkholderia*-like endobacteria (*Candidatus* Glomeribacter gigasporarum*; Ca*Gg) within its cytoplasm, which have been shown to impact fungal biology ^40,41,42^. The presence of *Ca*Gg in *G. margarita* spores (+*Ca*Gg) and the availability of a cured line (-*Ca*Gg), along with its large and repeated genome and phylogenetic placement, collectively position *G. margarita* as an ideal species to elucidate diversity and conservation of AMF chromosome biology and the malleability of their A/B compartments and TADs.

## Results

### Chromosome-level assembly, annotation, and phylogenomics of *G. margarita*

We performed PacBio and Hi-C sequencing of genomic DNA extracted from *G. margarita* +*Ca*Gg spores. Using Hifiasm with Hi-C integration mode ^43^, we assembled the HiFi long reads into draft contigs, which were then scaffolded using Hi-C data ^18,43^. For *G. margarita*, this approach generated 43 chromosome-level scaffolds, with a genome coverage of 82X, a size of 792.14 Mb and an N_50_ of 18.89 Mb (**Table 1**; **Fig. 1a, b**). Of these, 20 chromosomes have telomeres at both ends, 20 have telomeres at only one end, and three lack telomeres entirely. Only 23 contigs (0.718 Mb) could not be assigned to any chromosome. This assembly represents a significant improvement over the previously available datasets^39^ in assembly size (792.14 Mb vs. 773.10 Mb), fragmentation (43 vs. 6490 scaffolds), contiguity (N_50_ of 18.89 Mb vs. 326.79 kb), and gene count (30211 vs. 26603).

**Table 1.** Summary statistics for the genome assembly of *G. margarita*

| Feature | *G. margarita* |
| --- | --- |
| Genome size (Mbp) | 792.14 |
| No. of scaffolds | 43 |
| No. of genes | 30,211 |
| No. of rDNA clusters | 21 |
| Repeat content (%) (after curation) | 78.28 |
| Busco completeness (%) | 95.4 |
| GC (%) | 27.77 |

**Figure 1.**
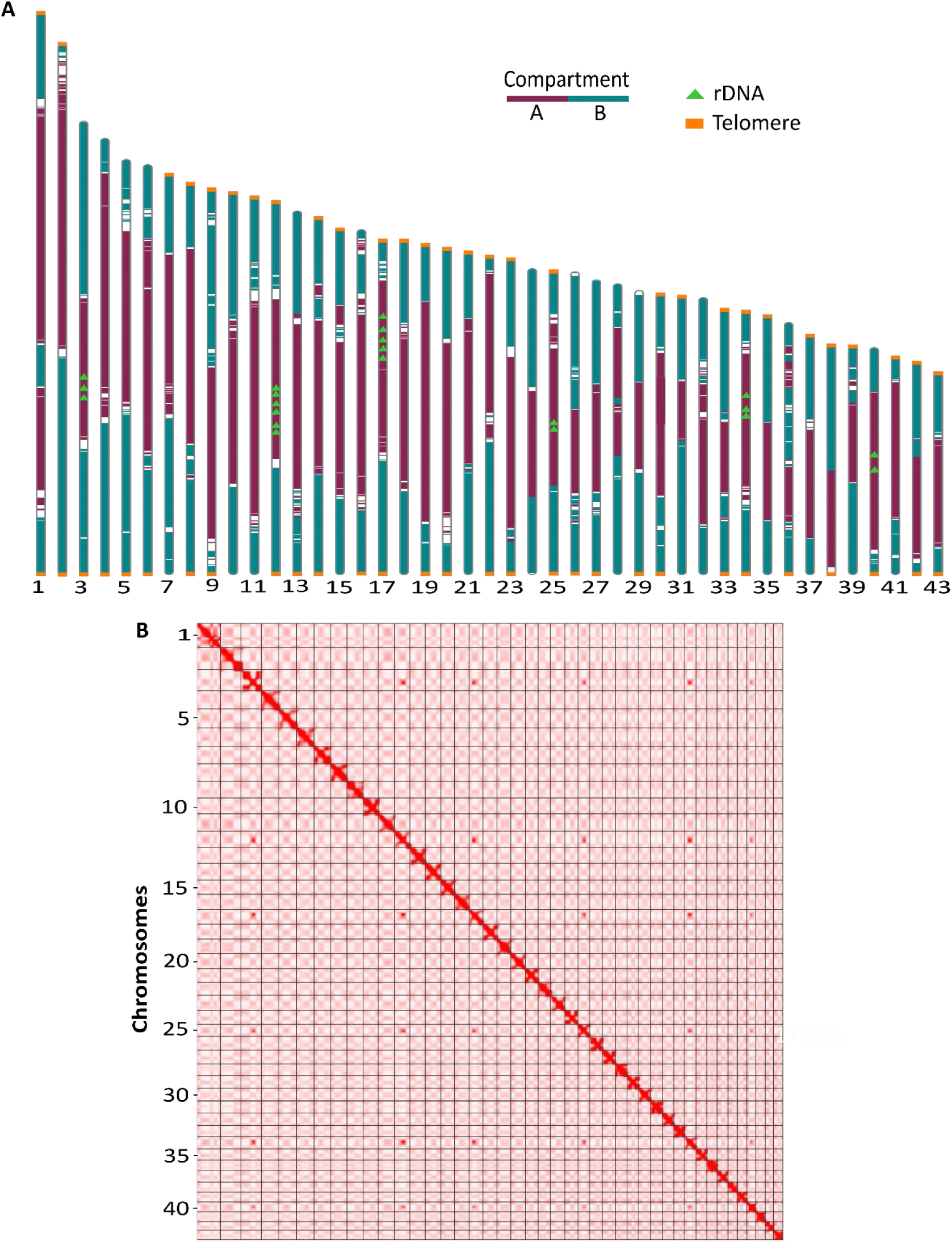
*Gigaspora margarita* chromosomes and genome content. (A) Karyoplots of 43 chromosomes, illustrating rDNA genes in lime green colour and A/B compartments in violet-red and turquoise colours, respectively, within the ideograms. (B) Genome-wide Hi-C contact map of *G. margarita*. The black squares represent chromosomes sorted by size. The colour intensity corresponds to interaction frequencies between loci, with darker red indicating high contact probability and white signifying low or no interactions.

Genome annotation identified 30,211 protein-coding genes, resulting in a BUSCO completeness of 95.4% (**Table 1**). Of these, 2,438 (8.1%) were annotated as putative secreted proteins and effectors, numbers higher than those reported in *R. irregularis* **(Table S2)**. We confirm that *G. margarita* lacks the hallmark “Missing Glomeromycota Core Genes (MGCGs)” ^44^ (**Table S3)** and has a reduced set of carbohydrate-active enzymes (CAZymes), but shows a significant expansion of CAZyme families involved in chitin metabolism (GH18, GT1, GT2, AA7, and CE4)^39,45^ (**Table S4)**. The annotation also uncovered cobalamin-dependent enzymes in *G. margarita* **(Table S5)**, supporting the hypothesis that the fungus uses cobalamin supplied by *Ca*Gg ^46^. The chromosome-level assembly confirms that all AMF with sequenced genomes carry all known meiosis-specific genes, and supports that, as opposed to most AMF with sequenced genomes^21^, including representatives of early branching lineages^47,48^, the genes composing the putative AMF mating-type^17,18^, namely a choline transporter, two homeodomain proteins (HD1-2), and a phosphate glycerate mutase, are not adjacent within members of the Gigasporaceae ^16^.

Alongside the nuclear genome, we recovered the complete mitochondrial genome as a single contig. The genome size is 96,986 bp, and it encodes core mitochondrial genes, including 19 protein-coding genes, 2 trans-spliced rRNA genes, and 24 tRNA genes, supporting previous findings of the *G. margarita* mitochondrion^53^.

We took advantage of this new chromosome-level dataset to determine the evolutionary placement of *G. margarita* relative to other AMF species via phylogenomic analyses (**Fig. S1**). These analyses support the sister relationship between Glomeromycotina and Mortierellomycotina within the Mucoromycota phylum ^1,49^. It also backs the close evolutionary relationship between Glomerales and Diversisporales, the paraphyletic placement of *Entrophospora* species ^45,48,50,51^, as well as the early branching of Paraglomerales and Archeosporales within the AMF phylogeny. Notably, in support of phylogenetic analyses based on ribosomal genes ^52^ and available genome datasets ^47^, our findings highlight Paraglomerales as representing the earliest AMF phylogenetic node. However, we were unable to fully reject an alternative placement of *P. occultum* with members of the Archaeosporales.

### The divergent rDNA operons are physically linked in *G. margarita*

The chromosome-level genome annotation also confirms that, unlike most eukaryotes, AMF carry few copies of highly divergent ribosomal DNA (rDNA) operons within their genomes. The *G. margarita* genome has 21 rDNA copies, approximately twice as many *R. irregularis* strains^24^, and these vary in copy number and sequence both within and across chromosomes 3, 12, 17, 25, 34 and 40 (**Fig. 1a, b**; **Table S6**). The high rDNA sequence paralogy within members of Glomerales was reported to cause significant taxonomic challenges, as rDNA paralogs can sometimes cluster across species boundaries ^54,55^. We find that this issue is exacerbated in *Gigaspora* species, further building concerns about the utility of these genes alone for AMF taxonomy ^54–56^. For instance, the rDNA paralogs from the *G. margarita* genome scatter across three clades and are shared with sequences from *G. decipiens*, *G. albida*, and *G. gigantea*. Almost all *Gigaspora* species show similar paraphyletic clustering based on rDNA paralogy rather than speciation (**Fig. S2**).

In addition to enabling chromosome-level assemblies, the Hi-C data also revealed strong physical interactions among rDNA copy regions on chromosomes 3, 12, 17, 25, 34, and 40, as evidenced by visually bright signals off the diagonal line in Hi-C maps (**Fig. 1b, Table S6**). In model eukaryotes, active rDNA copies physically cluster in the nucleolus to allow rDNA biogenesis, while inactive copies are found at its edges or outside ^57–59^. A very similar pattern is seen in *G. margarita*: most rDNA units are hypomethylated (i.e., actively transcribed), with only one unit, located on chromosome 12, highly methylated, and thus possibly representing a pseudogene. Notably, investigations of Hi-C data from *R. irregularis* strains uncovered similar rDNA interactions (**Fig. S3**) between regions on chromosomes 9, 18, 23, and 28. Taken together, our results indicate that AMF divergent rDNAs follow a cellular mechanism found in model eukaryotes, in which active rDNA units are physically close to each other within the nucleus, presumably residing within the nucleolus, while inactive units are positioned at the periphery of the nucleolus or outside it **(Fig. 2)**.

**Figure 2.**
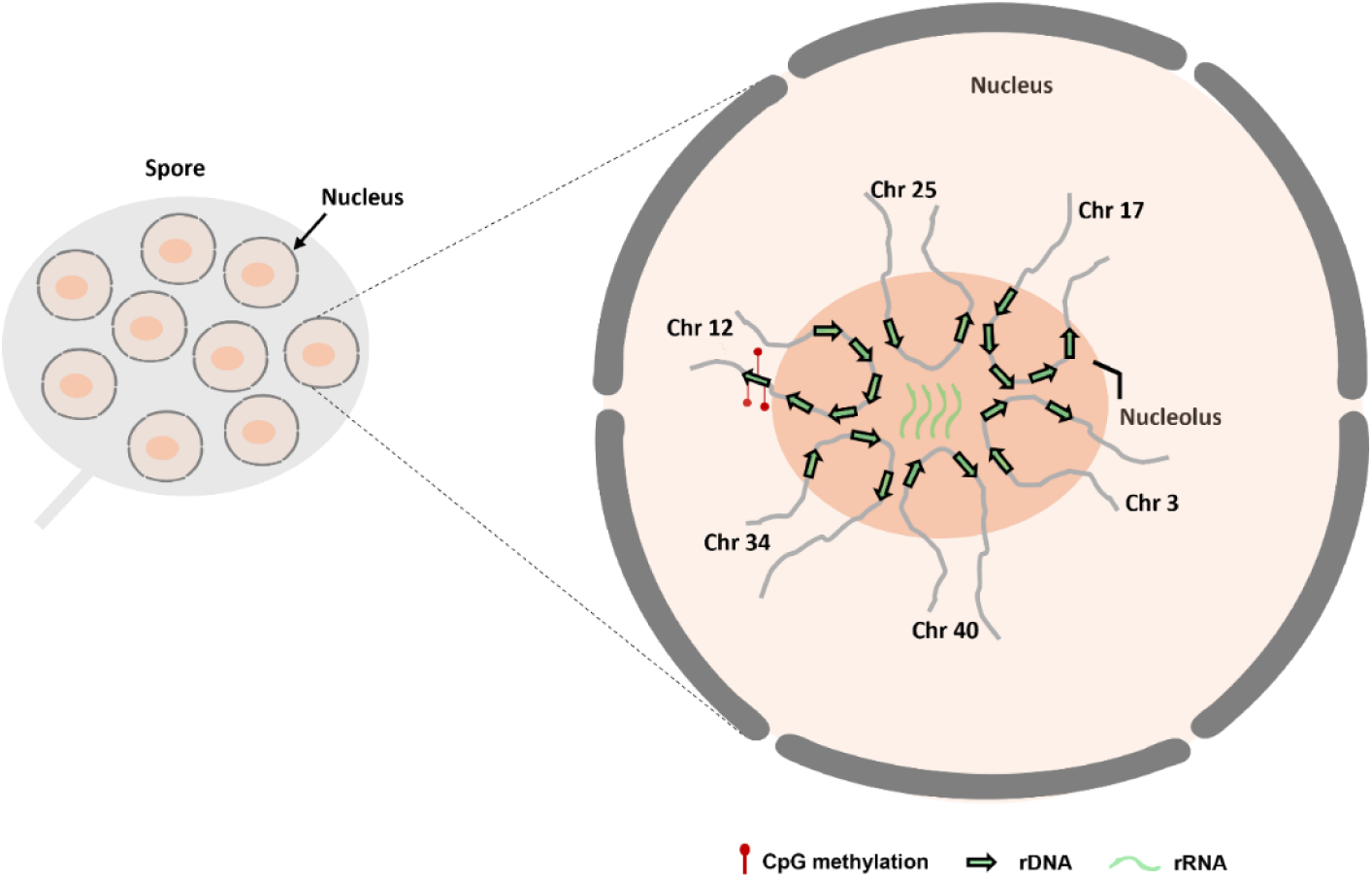
Model of the rDNA 3D organization within the AMF nucleolus. Based on current Hi-C data and knowledge of rRNA transcription in model eukaryotes. Transcriptionally active rDNA repeats from different scaffolds hypothetically interact physically in the nucleolus. The methylated rDNA copy sits outside the nucleolus.

### Endobacteria-driven regulation of transposable elements in *G. margarita*

The genome of *G. margarita* is known to be highly repetitive, but the true extent and diversity of these repeats was likely obscured by the fragmented nature of available genome datasets ^60^ and a lack of curated analysis of such repeats ^25^. Curating our chromosome-level assembly revealed that 78.34% of the *G. margarita* genome consists of repeats, of which 56.1% belong to known TE families. Among these, 19.2% correspond to DNA/TIRS elements (i.e., DNA transposons), while 17.1% and 13.1% are LTR and LINE elements, respectively. Overall, 22.7% of the repeats remain unknown and cannot be classified after manual curation (**Fig. 3**), which compares to 41% in previous work ^39^. These unknown repeats may represent expanded gene families commonly found in AMF^25,61^, as well as TE families that have not yet been formally characterized in model species, including fungi. We assessed the expansion and degeneration of TEs in *G. margarita* by constructing their repeat landscapes based on Kimura substitution calculations (**Fig. 3**), where lower Kimura substitution rates indicate recent TE insertions, while higher rates suggest older insertions ^61^. Our analyses build on previous findings showing that TEs in Diversisporales are enriched in recent and active expansions ^25^, with DNA/TIR elements, LINEs, and LTRs being the primary contributors to recent TE bursts in *G. margarita*.

**Figure 3.**
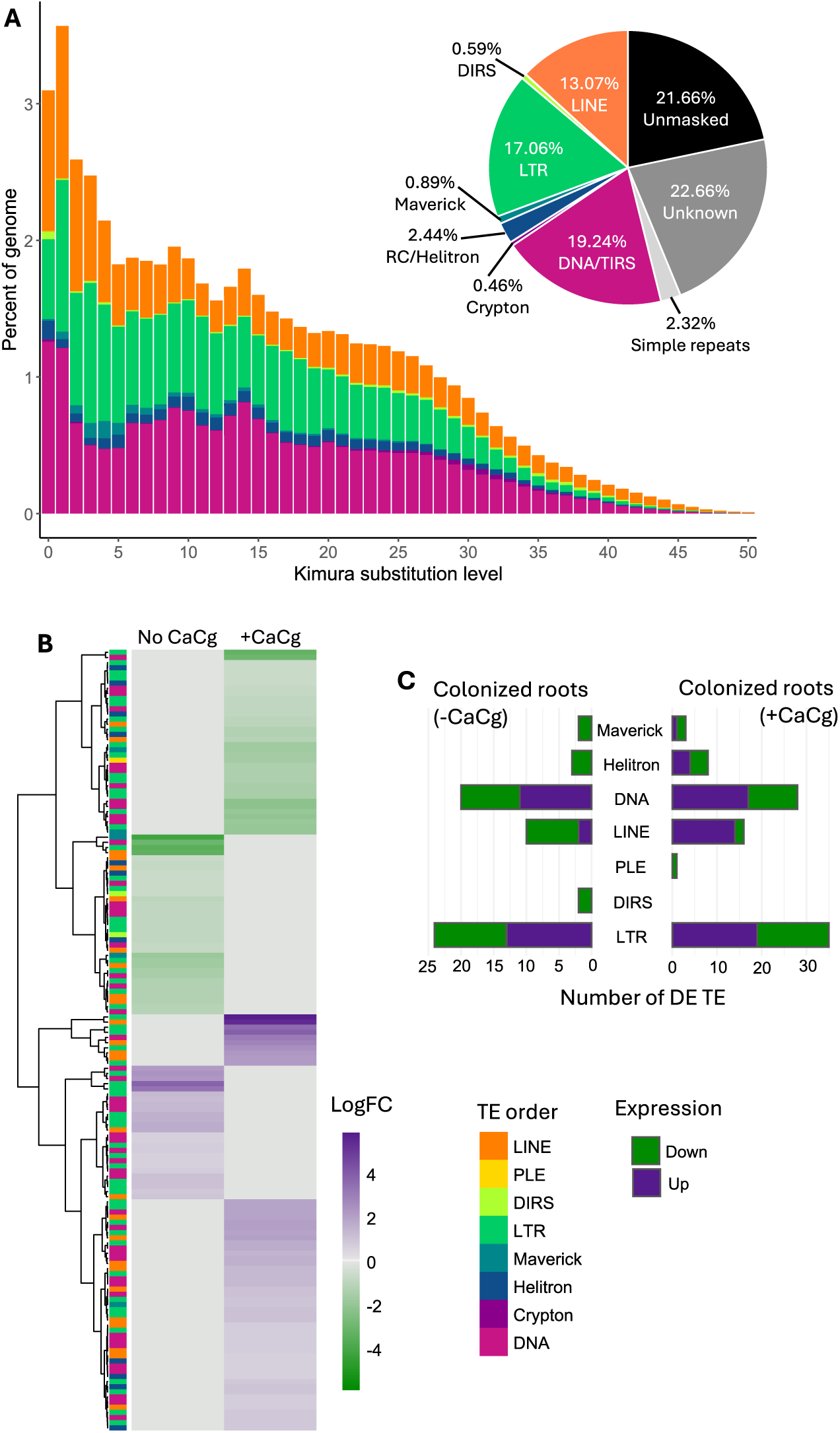
Composition and activity of transposable elements in *G. margarita* genome. (A) Transposable element distribution and repeat landscape. The pie chart depicts the relative abundance of each type of sequence in the *G. margarita* genome. The histogram represents the repeat landscape grouped by Kimura divergence levels (x-axis) in relation to the genome (x-axis). (B) Heatmap for differentially expressed TE families during *Lotus japonicus* root colonization, comparing samples lacking the *Ca*Cg endobacterium (-*Ca*Cg) with *Ca*Cg-containing samples (+*Ca*Cg) in relation to spores (control). Rows represent TE families, annotated by TE order and clustered by expression profile. (C) Number of differentially expressed TE families by order and direction of regulation (up- or down-regulated) in colonized roots, shown separately for samples without and with *Ca*Cg.

In AMF, including *G. margarita*, TEs are preferentially located close to promoters and genes involved in molecular communication with the host (e.g., effectors) ^25^, which indicates their potential role in regulating gene expression and maintaining successful mycorrhizal relationships. Reports of significant TE upregulation in the model AMF *R. irregularis* following root colonization supported this view ^27^. A more comprehensive annotation of TEs allowed a detailed analysis of the differential regulation of fungal TEs in *Lotus japonicus* roots colonized by *G. margarita*, with and without endobacteria. Remarkably, the presence of the bacterial endosymbiont alters the regulation of specific TE families during root colonization (**Fig. 3b, c**). Among these, some members of the Helitron, Maverick, and LINE families were specifically upregulated only in the presence of *Ca*Cg, while members of the PLE family were upregulated exclusively in the absence of *Ca*Cg. Furthermore, DIRS families were downregulated in the absence of *Ca*Cg (**Fig. 3c**).

### Identification of A/B compartments and Topologically Associating Domains in *G. margarita*

Hi-C data also revealed that the *G. margarita* genome exhibits a checkered pattern delineating two A/B compartments **(Figure 4a, b; Fig. S4)**, as reported in all *R. irregularis* strains with available Hi-C data ^18,24^. Some chromosomes carry large blocks of A or B compartments **(Fig 4a)**, while others harbour a finely interleaved, plaid-like pattern of A/B compartmentalization (**Fig. 4b**).

**Figure 4.**
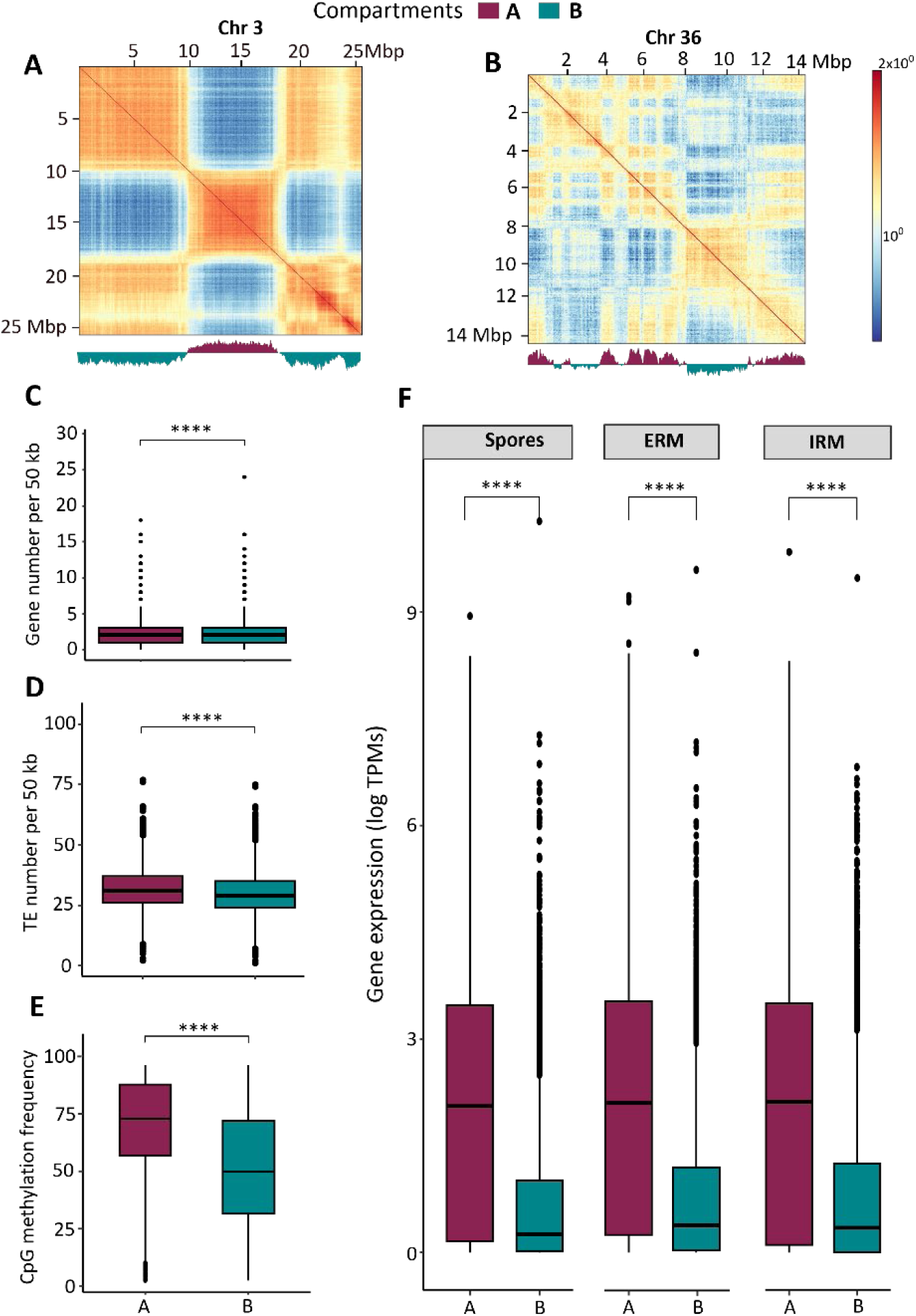
Genome compartmentalization in *G. margarita.* Examples of Hi-C contact maps showing A/B compartments in *G. margarita* chromosomes (A) 3 and (B) 36. The bottom track shows the first eigenvector values, identifying the compartment interaction at 50-kb resolution. Regions that interact more frequently are visualized as brighter squares on the contact map. Gene/repeat densities, methylation frequency and gene expression in compartments A and B of *G. margarita*. Boxplots showing (C) Genes (Genes per 50 kb), (D) repeat densities (Repeats per 50 kb), (E) CpG methylation frequency, and (F) gene expression levels (logTPM + 1) in three conditions: Spores, ERM, and IRM (*L. japonicus*) in A and B compartments. Boxes show the first quartile (25%), the middle black line (50%), and the third quartile (75%). The whiskers extend to 1.5× the box length, representing the outliers as dots. Asterisks above the boxplots indicate significant differences between the A/B compartments (*p* < 0.05, Wilcoxon rank-sum test).

The *G. margarita* A compartment has significantly higher gene density and average gene expression compared to the B compartment and contains most core genes and all rDNA operons (**Fig. 4a**), while the B compartment is significantly enriched for secreted proteins and candidate effector genes (**Table S7**). Although both compartments have similar TE densities **(Fig. 4d)**, the TEs in the A compartment are significantly more methylated **(Fig. 4e)**. Like *R. irregularis* homokaryons^24,61^, *G. margarita* shows a strong bimodal distribution of methylation levels, with a larger percentage of CpG sites either highly methylated (>8 = 58.8%) or weakly methylated (<2 = 38.6%). Notably, A/B compartments remain remarkably stable between the +*Ca*Cg and -*Ca*Cg conditions – i.e., no significant change in checkered patterns was observed both within and among chromosomes. In *G. margarita*, the A compartments are preferentially located within chromosome cores (Fig 1), while B-compartments are generally found at the chromosome ends. This clear distinction in A/B localization within chromosomes is not present in *R. irregularis* isolates ^24^.

Hi-C data analysis also revealed 1,407 TAD-like structures in *G. margarita* (**Fig. 5a**). The TADs cover 92.16% of the genome, with a median size of 420 kb (**Fig. 5b**), in line with reports from other eukaryotes, including non-AMF fungi ^30,32,62^.

**Figure 5.**
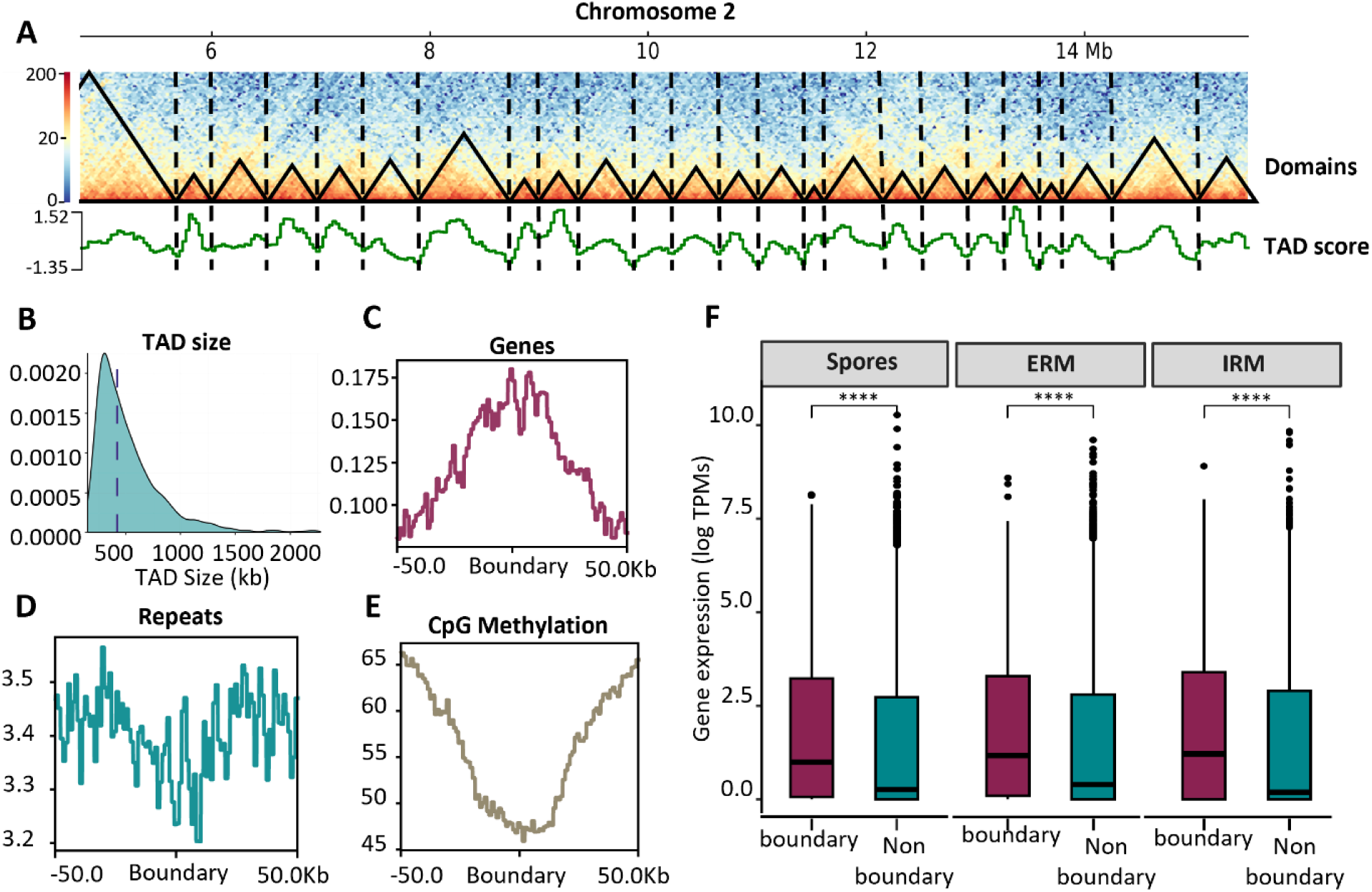
Topologically associating domains (TADs)-like structures in the *G. margarita* genome. (A) A representative section of chromosome 2 showing examples of TADs as black triangles. The bottom green line corresponds to the insulation score (TAD score), and the vertical black dashed lines highlight the predicted TAD. (B) Density plot showing TAD size distribution. The vertical dashed line represents the mean value. (C-E) Line plots of (C) genes, (D) repeats, and (E) DNA methylation (CpG) at domain boundaries and ±50 kb from boundaries. (F) Gene expression (log TPMs + 1) in boundary and non-boundary regions across three conditions: Spores, ERM, and IRM (*L. japonicus*), respectively. Boxes show the first quartile (25%), the middle black line (50%), and the third quartile (75%). The whiskers extend to 1.5× the box length, representing the outliers as dots. Asterisks above the boxplots indicate significant differences between the boundary and non-boundary regions (*p* < 0.05, Wilcoxon rank-sum test).

We identified TAD boundaries based on low insulation scores and, surprisingly, found that they are gene-rich and depleted of repeats (**Fig. 5c, d**). The gene enrichment at boundary regions is linked to low methylation levels (**Fig. 5e**) and, accordingly, boundary-associated genes have significantly higher expression levels across multiple life stages, including germinating spores, extraradical, and intraradical mycelium (*L. japonicus*), compared to non-TAD boundaries (**Fig. 5f**). Moreover, genes within the same TAD are co-expressed across life stages (**Fig. S5**), indicating that TAD boundaries function as transcriptional hotspots in AMF.

### The tripartite *Ca*Gg endosymbiotic genome supports genomic plasticity and reveals novel pathways for enhanced stress defence and cofactor biosynthesis

Alongside the AMF genome, we obtained the complete genome of the *Ca*Gg endosymbiont, comprising a single circular chromosome of 1,998,997 bp and two circular plasmids, referred to herein as p*Ca*Gg01 (99,883 bp) and p*Ca*Gg02 (22,198 bp), with a read depth double that of the chromosome **(Fig. 6; Table 2)**. The endobacterial circular chromosome and plasmid assemblies contain the expected ORI motifs and were all corroborated by uniform read coverage, as well as Hi-C contact mapping (**Fig. S6**) and independent assemblers ^63,64^.

**Figure 6.**
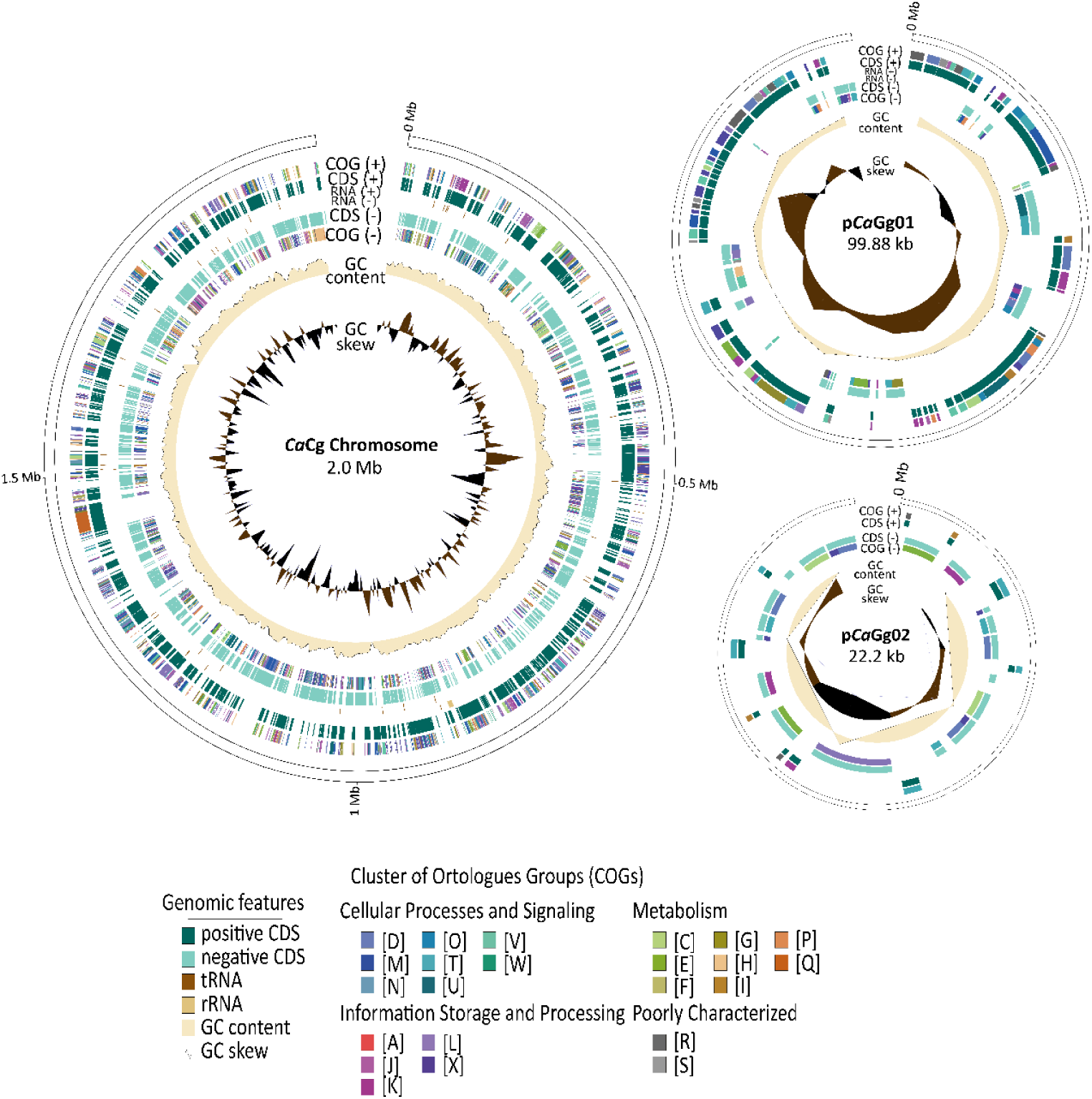
Circular genome map of *Ca*Gg. From outside to inside: + strand clusters of orthologous genes (COGs); coding sequences (CDS) on the + strand; rRNA and tRNA on the + strand; CDSs, rRNA, and tRNA on the – strand; COGs on the - strand; GC content; GC skew.

**Table 2.**
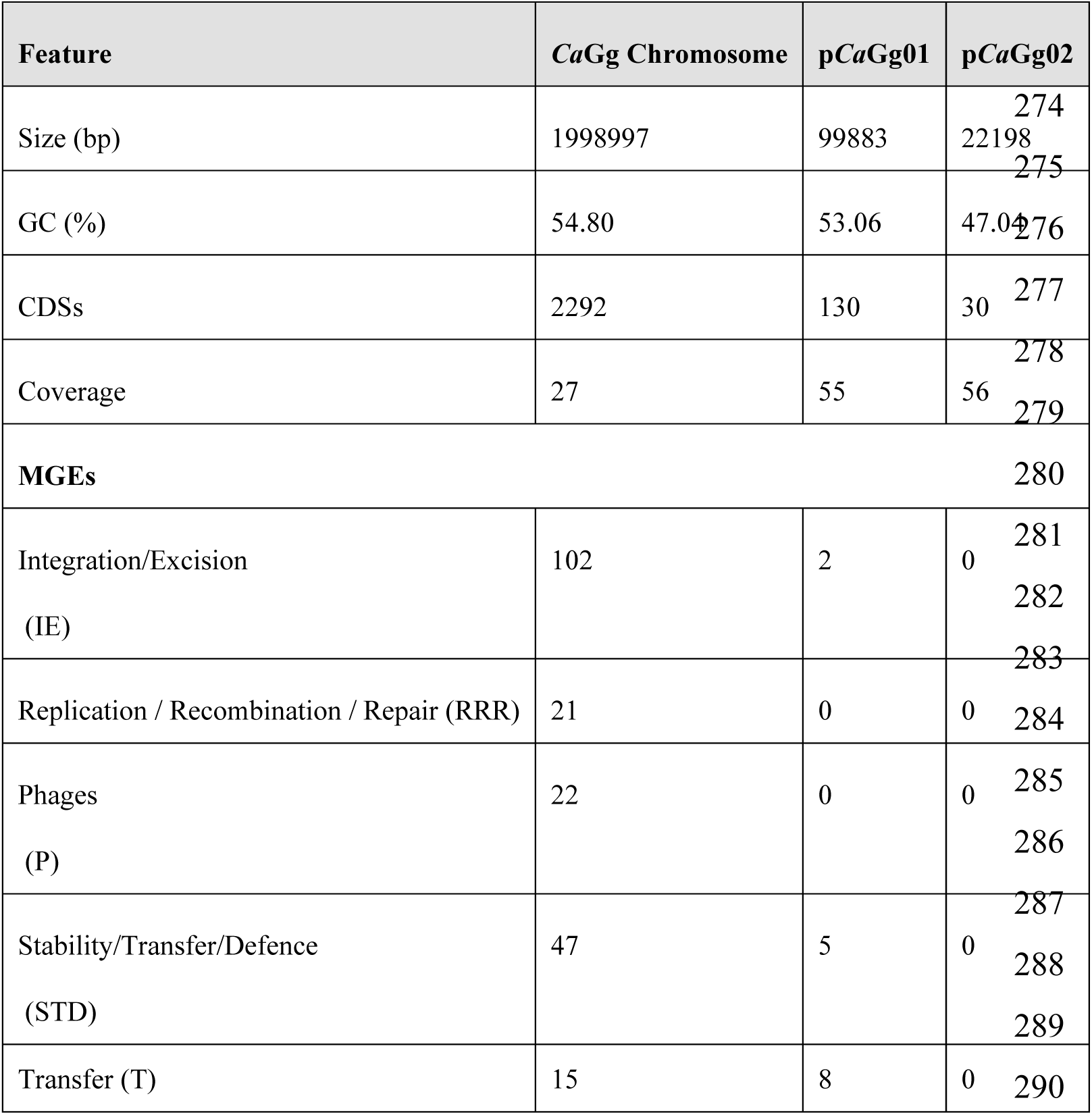
Genome statistics and mobile genetic elements (MGEs) of the *Ca*Gg chromosome and associated plasmids.

This new assembly recovers substantially more coding capacity and non-coding features than the previously fragmented version did. Specifically, our genome assembly, totalling 2,121,078 bp, is 20% larger than the previous fragmented assemblies obtained with pyrosequencing and fosmid libraries, which reported 3 ORIs in 4 genomes at 15x coverage across 125 contigs ^46^. It is also complete, fully resolving previously undetected plasmid sequences. The *Ca*Gg chromosome contains 2,292 protein-coding genes, 44 tRNAs, 21 non-coding RNAs, one transfer-messenger RNA, and a single complete rRNA operon. The plasmid p*Ca*Gg01 has 130 coding genes and one ncRNA, while p*Ca*Gg02 has 30 coding genes. A cluster of Orthologous Genes (COGs) analysis reveals that the *Ca*Gg chromosome contains a high proportion of genes associated with mobilome (X), while p*Ca*Gg01 is dominated by genes classified as defence (V), and p*Ca*Gg02 mainly contains genes involved in signal transduction (T) (**Fig. 6**).

The metabolic reconstruction of the *Ca*Gg chromosome confirms previously reported core metabolic modules, including the absence of phosphofructokinase (*pfk*), fatty acid degradation genes, reduced amino acid biosynthesis enzymes, and a complete cobalamin (B12) biosynthesis pathway. The endosymbiont also encodes multiple membrane transport systems, including ABC transporters and members of the major facilitator superfamily (MFS), as well as Type II, III, and IV secretion systems and complete Sec and Tat pathways ^46^.

Our improved assembly also uncovered novel and biologically relevant metabolic capabilities of *CaGg*. These include complete pathways for coenzyme A, NAD, lipoic acid, PreQ1, and heme metabolism, revealing a much broader capacity for cofactor production than previously assumed. We also found a CRISPR array lacking a cas-motif. This orphan array may be linked to the identification of phage elements in our assembly, including two intact prophages and several remnants on the *Ca*Gg chromosome **(Table S8)**, thereby explaining the remarkable enrichment of Mobile Genetic Elements (MGEs) **(Table 2)**. Remarkably, some of the MGEs are classified as plasmid-origin gene families, and upon further examination, we identified several plasmid-related genes in the *CaGg* genome, including some involved in plasmid replication, partitioning, stabilization, and conjugation, altogether suggesting possible distinct plasmid integrations into the *CaGg* genome **(Table S9)**. None of these putative insertions result in coverage drops or changes in Hi-C signals, indicating that these are not artefactual.

Overall, the MGEs cover 6.97% of the *CaGg* genome and are classified into five functional categories: Integration/Excision (IE, 102), Replication/Recombination/Repair (RRR, 21), Phage (P, 22), Stability/Transfer/Defense (STD, 47), and Transfer (T, 15). As observed in other Glomeromycotina endosymbionts ^65^, IE elements are enriched in the genome, suggesting that they are major contributors to genome plasticity in Glomeromycotina endosymbionts. The high mobilome content prompted us to analyze type II toxin-antitoxin (TAs) modules, which are thought to stabilize MGEs ^66^ and mediate stress-induced persistence in *Ca*Gg ^67^. The present assembly uncovered 41 new TAs (39 chromosomal and 8 on p*Ca*Gg01), up from 9 in earlier predictions ^67^ (**Table S10**).

## Discussion

In this work, we reported the first chromosome-level assembly and 3D analysis of an AMF genome outside the Glomerales. This provided an unprecedented view into the biology of one of the largest and most repeat-rich genomes known within the fungal kingdom and uncovered novel insights into the biology and evolution of its bacterial endosymbiont.

### A chromosomal view of a large and highly repetitive non-model AMF

Combining long reads with chromatin-capture methods has successfully helped assemble the large, highly repetitive *G. margarita* genome and its endosymbiont into a complete chromosome-level assembly. This allowed us to demonstrate that AMF have more chromosomes than most other fungal relatives and that, with 43 chromosome-level scaffolds, *G. margarita* likely has the highest reported chromosome count to date in the fungal kingdom. Our chromosome annotation reveals that *G. margarita* encodes approximately 30,211 genes and retains all the hallmarks of obligate biotrophy typical of AMF ^48^. These include a reduced set of cell wall-degrading enzymes, the lack of fatty acid biosynthesis genes, the loss of thiamine synthesis genes, and the uptake of soluble sugars, such as glucose, which are essential compounds they obtain directly from their hosts ^12,16^. We also report a larger-than-expected set of proteins potentially involved in the dialogue with the host ^24,26,68,69^, suggesting an improved ability to establish symbioses with multiple plants, and we found primary evidence that *G. margarita* utilizes cobalamin produced by its endobacterial host, *Ca*Gg.

Beyond gene content, the chromosome-level assembly provides a curated and comprehensive view of TE diversity in a genome long recognized for its repetitiveness and fragmentation ^39^. While previous work linked the large genome size of *G. margarita* primarily to LINE expansions ^39^, our refined TE annotation uncovers a substantial contribution of Class II elements, including DNA/TIR transposons and Mavericks in *G. margarita* genome biology. These DNA transposons have been implicated in gene duplication, genome restructuring, and regulatory innovation in fungi and other eukaryotes ^60^, suggesting that multiple TE classes, not only retrotransposons, continue to contribute to genome complexity in *G. margarita*. Our analyses also continue to support the view that, in addition to dictating genome evolution, the AMF TEs are major players in regulating gene expression during colonization, presumably due to their localization within regulatory regions and in proximity to candidate secreted proteins and effectors. It is noteworthy that different TEs families are regulated in *G. margarita* and in *R. irregularis* strains during root colonization. This could mirror lineage-specific regulatory adaptations in AMF and/or host plant-driven TE regulations^27^ . It will be important to see how these elements are regulated across additional hosts and conditions in future studies, particularly in the presence or absence of the endobacterium.

The genome analyses of a large AMF genome also reinforce the growing evidence that rDNA sequence diversity is significant within individual AMF genomes. This continues to highlight the challenges of using this locus alone for taxonomic resolution, as divergent rDNA copies cluster by paralogy rather than by speciation, and supports the need for population genetics-based approaches in AMF taxonomy ^54,70^. Our evidence that some rDNA copies are likely pseudogenes further exacerbates these challenges.

### Remarkable conservation in genome biology among AMF lineages

In addition to sharing losses in key genes associated with obligate biotrophy and high rDNA gene paralogy, the genome biology of *G. margarita* follows patterns conserved among distinct AMF lineages. All species investigated to date with Hi-C have genomes partitioned into an A compartment with high gene density, expression, and repeat methylation, and a B compartment with low repeat methylation and an enrichment in secreted and effector proteins. This spatial organization of AMF chromatin likely maintains genome stability by silencing repeats in the A compartment, which is rich in housekeeping genes ^24^ while allowing the B compartment to serve as a hotspot for diversifying secreted proteins and effector encoding genes. Altogether, these features continue to underpin striking analogies in the genome biology and evolution of AMF and filamentous plant pathogens ^71,72^.

Despite the shared similarities, AMF genomes exhibit some notable epigenetic distinctions. For example, while the *R. irregularis* heterokaryons show a tripartite genome-wide CpG methylation distribution ^18^, the homokaryons and *G. margarita* display a bimodal methylation pattern ^24,73^. The reasons for these differences remain unknown, although it is tempting to speculate that the nuclear type (homokaryon vs. heterokaryon) influences AMF methylation distribution patterns. Our work also identified notable distinctions between TADs in AMF and those of fungal relatives. For example, in the fungal species *Epichloë festucae*, AT-rich, repeated blocks contribute to the formation of TADs ^35^, yet the repeat-depletion at the boundaries we observed suggests an alternative mechanism for TAD establishment in AMF that is more consistent with reports from other eukaryotic lineages^33,34,62,74^, whereby boundary genes are co-expressed across different conditions, indicating that TAD organization facilitates transcriptional coordination in these prominent symbionts.

### A model that explains the co-transcription of divergent rDNA operons in AMF

In most eukaryotes, rDNA genes exist as hundreds to thousands of identical copies scattered across different chromosomes, yet these copies are all transcribed in a spatially coordinated manner within the nucleolus ^57,58,75^. The localization of rDNA genes within the nucleolus has been primarily studied using microscopy techniques, such as fluorescence in situ hybridization (FISH) and immunofluorescence ^57,76^. Recently, 3C-based methods, such as Hi-C, have been employed to identify these interactions at a genome-wide scale, providing an additional independent line of evidence for rDNA co-localization ^77^. In our work, the presence of fewer, divergent rDNA copies in *G. margarita* has likely facilitated their capture and separation by Hi-C, thereby enabling the detection of their strong physical interactions, indicating that these genes co-localize, presumably within the nucleolus, to ensure their co-transcription and enhance ribosome biogenesis^57,58^. In addition, the identification of identical signals in *R. irregularis* ^24^ shows that the co-localization of rDNA is a conserved mechanism of rDNA organization and co-regulation in AMF.

### An improved view of endosymbiotic contribution to *G. margarita*

Our complete genome assembly and comparative analysis revealed that the *Ca*Gg genome comprises a large circular chromosome and two smaller complete circular plasmids, each with distinct ORI motifs, supported by HiFi and Hi-C data and independent assemblers. As such, this work clarifies previous assumptions, which suggested that the *Ca*Gg genome is organized into 2 to 4 genomic units ranging between 1.4 to 2.5 in size ^46,78^. Like other AMF endosymbionts ^40,79,80^, the *Ca*Gg genome is smaller than that of free-living *Burkholderia* relatives^81^, resulting in limited metabolic capabilities, including the inability to utilize glycolysis as an energy source and a limited capacity to biosynthesize essential amino acids, underscoring the AMF host’s strong dependence.

The close-knit relationship between *Ca*Gg and *G. margarita* is further emphasized by our finding that some fungal TEs are upregulated only in the presence of the endobacterium, and the identification of novel pathways through which *Ca*Gg may influence the host’s fungal physiology. In particular, the presence of a complete heme biosynthesis pathway suggests an unsuspected role in supporting AMF host bioenergetics, consistent with observations in other host-endosymbiont systems ^82^. Indeed, heme is an essential cofactor for many proteins, including cytochromes of the mitochondrial electron transport chain, which are highly abundant in the *G. margarita* genomes. As such, it is intriguing to speculate that *Ca*Gg-derived heme enhances the activity of respiratory complexes in the AMF host, providing a possible explanation for the higher ATP production and respiratory activity reported in endobacteria-containing *G. margarita* compared to the cured line ^83^. In other filamentous fungi^84^, heme also plays roles in growth, development, and stress adaptation, suggesting that *Ca*Gg-encoded heme production could affect multiple aspects of AMF physiology beyond respiration. Future analyses will hopefully reveal how these novel endobacterial genes are regulated in response to environmental changes.

Another surprising discovery was that *Ca*Gg carries several MGEs, including prophages, expanding the viral pool within the *G. margarita* symbiotic system ^41^. While most bacteria rely on CRISPR-Cas systems to defend against foreign DNA^85,86^, *Ca*Gg, like many endosymbionts, lacks a functional CRISPR-Cas system ^87,88^. The presence of this orphan CRISPR array likely explains the high MGE content, including prophages, in the *Ca*Gg genome, and it indicates that CRISPR-based defence was active in the BRE endobacterial ancestors prior to some lineages becoming obligate endosymbionts. Conversely, the enrichment of toxin-antitoxin systems in *Ca*Gg might represent an alternative defence strategy in the absence of CRISPR-Cas ^86,89^.

## Conclusions

This study presents the first chromosome-scale and 3D genome assembly of a non-model AMF, revealing conserved principles of chromatin architecture among AMF despite dramatic differences in genome size and repeat content. Our findings confirm that A/B compartmentalization and TAD-like structures are fundamental features of AMF genome organization, supporting coordinated gene expression and genome stability. We also uncover a novel mechanism for rDNA co-localization within the nucleolus, which likely facilitates ribosome biogenesis despite extensive rDNA sequence divergence—a hallmark of AMF genomes.

Beyond fungal chromatin biology, our work sheds light on new functional contributions of the obligate endobacterium *Ca*Gg by revealing a multipartite architecture enriched in mobile genetic elements, underscoring its genomic plasticity that cannot be offset by an incomplete CRISPR-Cas system. Additionally, we identify novel pathways for essential cofactors, including heme and cobalamin, suggesting a direct role in enhancing host bioenergetics and stress resilience. The observation that *Ca*Gg presence modulates transposable element activity in *G. margarita* further highlights a complex interplay between host and symbiont at the epigenetic and genome stability levels.

Together, these findings point to a model in which chromatin architecture, rDNA organization, and endosymbiont-derived metabolic capabilities collectively shape the biology and ecological success of AMF. Future work should explore how these interactions respond to environmental cues and influence plant fitness, including in early-branching lineages, to provide complete insights into the molecular foundations of one of the most impactful plant symbioses.

## Methods

### Sample preparation for HiFi sequencing

Spores of *Gigaspora margarita* Becker and Hall (BEG 34, deposited at the European Bank of Glomeromycota) containing (+*Ca*Gg) or lacking (-*Ca*Gg) the obligate endobacterium *Candidatus* Glomeribacter gigasporarum were used in this study. The -*Ca*Gg spores were obtained from +*Ca*Gg spores as described in ^90^. The +*Ca*Gg and -*Ca*Gg spores were isolated from clover trap plants by wet sieving^91^ and collected individually under a dissecting microscope. Only the healthiest spores were selected and pooled into groups of 100, then surface-sterilized with a solution of Chloramine T (3% w/v) and streptomycin sulphate (0.03% w/v) in sterile distilled water. A batch of +*Ca*Gg spores was immediately frozen in liquid nitrogen, while the remaining batches (+*Ca*Gg and -*Ca*Gg) were incubated to allow germination.

### DNA extraction and purification

DNA was extracted from pools of 100 sterilized and frozen +*Ca*Gg spores using the DNeasy Plant Pro Kit (Qiagen). The spores were ground in the extraction buffer using a TissueLyser homogenizer (2 min at 24 Hz, repeated twice), and DNA extraction was performed according to the manufacturer’s instructions, except that the PS solution was omitted and the samples were not heated. DNA from two independent pools of 100 spores each was combined, and a cleanup step was performed with the DNeasy PowerClean Pro Kit (Qiagen) according to the manufacturer’s instructions.

### Sample preparation and Hi-C sequencing

For Hi-C analysis, 2,900 +*Ca*Gg and 2,900 -*Ca*Gg sterilized spores were placed in multiwell plates containing 1 mL of sterile distilled water and incubated in the dark at 30 °C for 7 days to allow germination. Germinated spores and pre-symbiotic mycelium were fixed at room temperature under gentle shaking for 20 min in 1% (v/v) formaldehyde in 0.01 M PBS (pH 7.2), followed by an additional 15 min incubation in 0.125 M glycine in 1% formaldehyde in 0.01 M PBS (pH 7.2). Spores and mycelium were then frozen in liquid nitrogen and ground with sterile micropestles to a fine powder. Samples were stored at −80 °C until library preparation was performed.

Hi-C data was processed using the Proximo Hi-C Kit (Fungal) from Phase Genomics (Seattle, WA, USA), employing four restriction enzymes: DpnII, MseI, DdeI, and HinFI. The Hi-C libraries were sequenced on the Illumina NovaSeq X PE150 platform, generating 194 and 202 million reads for the -*Ca*Gg and +*Ca*Gg samples, respectively **(Table S1)**.

### Genome assembly and Hi-C scaffolding

A total of 9.88 million PacBio HIFI reads were obtained on the Revio Platform at Mount Sinai Hospital (Toronto) and assembled into contigs using Hifiasm v0.16.1 with the parameters -l0 (no purging) and --h1 and --h2 for Hi-C data integration ^93^. The raw assembly was queried against the NCBI nr database using Diamond blastx (v0.9.14.115; perc_identity 75, -evalue 1e-5) to detect and remove contaminants, and identify mitochondria and *Ca*Gg contigs, which were annotated using MitoHifi v3.2.3 ^94^ and Bakta v1.8.2 ^95^, respectively.

The remaining contigs were scaffolded using the Hi-C reads, which were first processed using the Arima Genomics pipeline ^96^. The resulting BAM file alignments were used as input to the YaHS scaffolding tool ^97^. The assembled scaffolds were manually curated using PretextView ^98^ to generate chromosome-scale scaffolds and identify the unplaced contigs. A Python script was used to identify telomeres (https://github.com/Jana_Sperschneider/FindTelomeres). The final chromosome-scale scaffolds were visualized using the genome-wide Hi-C heatmap generated by Juicebox Assembly Tools v1.11.08^99^ and through a chromosome ideogram layout using RIdeogram^100^.

### Repeat masking and transposable elements analysis

The assembled genome was submitted to RepeatModeler2^101^ to generate a consensus library of repetitive sequences. To better classify repeats and address the high number of unclassified sequences (“Unknown”), we curated the consensus library using TEtrimmer^102^ and manual curation, as has been previously done for other AMF species ^25,27^. Using this method, sequences known to be expanded genes were removed. The genome masking and TEs annotation were performed on the final library, which contains a curated consensus of the assembled genome using RepeatMasker^103^.

RNA-seq data from intraradical mycelium with and without the endobacterium, from colonizing *Lotus japonicus* roots, and from extraradical mycelium of *G. margarita* (PRJNA267628)^104^ were used to assess the differential expression of TEs. The reads were first aligned to the reference genome assembled in this study using HISAT2. TE expression was then evaluated using TEtranscripts^105^, applied to the aligned reads, guided by the gene and TE annotations carried out in this study.

### Genome annotation

Gene prediction was performed on a soft-masked genome assembly using the Funannotate pipeline v1.8.15 ^106^. First, the command “funannotate train” was executed to train *ab initio* gene predictors with previously published RNA-seq data (PRJNA751155)^104^. Next, the command “funannotate predict” was run with the parameters --optimize_augustus and --ploidy 1. Multiple sources of evidence were utilized as input during the prediction of protein-coding sequences: (1) transcript assemblies (--transcript_evidence) and alignments (--rna_bam); (2) gene models generated by PASA (--pasa_gff); and (3) protein sequences from UniProtKB and the *G. margarita* proteome. The quality of the annotated genome was assessed using Benchmarking Universal Single-Copy Orthologs v5.2.2 (BUSCO) ^107^. SignalP v6.0 ^108^ was employed to predict secreted proteins with parameters --organism eukarya and --mode slow. EffectorP v3.0 ^109^ was utilized to predict effector genes, and carbohydrate-active enzymes (CAZymes) were annotated using dbCAN ^110,111^. To identify B12-dependent enzymes in *G. margarita*, homologs previously identified in *R. irregularis*^112^ were used as BLASTP queries.

### Phylogenetic analyses

To infer the phylogenetic relationships among SSU-ITS-LSU rDNA paralogs found in *G. margarita* and rDNA copies from other *Gigaspora* species, a dataset was created including sequences of *Dentiscutata savannicola* and *Fuscutata heterogama* as outgroups. Members of *Paradentiscutata* and *Intraornatospora*, the closest genera to *Gigaspora*, were not chosen as outgroups because they only have partial LSU sequences. Among the 21 rDNA operons of *G. margarita*, 19 were used as overlapping the partial SSU-ITS-LSU region used for phylogenetic inference. Overall, the dataset comprised 96 sequences representing eight *Gigaspora* species and 10 outgroup sequences. The dataset was aligned with the online version of MAFFT v.7^113^, using the E-INS-i iterative refinement method, and edited in MEGA v.5.2.2. by manual trimming of overarching and misaligned ends. Maximum likelihood phylogenetic inference was carried out in RAxML-NG via CIPRES Science Gateway 3.1^114^ and in IQ-TREE v.2.2.5 ^115^, with 1000 bootstrap, 1000 ultrafast bootstrap replicates and 1000 SH-aLRT tests, respectively. Additionally, a Bayesian analysis was performed in MrBayes v3.2.6 with 40 million generations and a stop rule at split frequency standard deviation = 0.01. All analyses were conducted with partitions and nucleotide substitution models as previously described ^116^. Notably, we found that most ingroup branches were either unsupported or poorly supported, particularly with respect to bootstrap and posterior probability values. In the Bayesian analysis, after 40 million generations, the average standard deviation of split frequencies did not approach 0.01 but instead fluctuated around 0.055, suggesting a potential lack of convergence in some bipartitions.

To determine the placement of *G. margarita* in the AMF phylogeny, phylogenomic analysis was performed using the “fungi_odb10” dataset from the BUSCO v4.0 ^107^. Profile-Hidden-Markov-Models corresponding to these markers were used to identify homologous sequences in *G. margarita* and 50 additional fungal genomes, using HMMER3 v3.1b2 as implemented in the PHYling pipeline (https://zenodo.org/records/10129968). A total of 702 conserved, single-copy orthologous proteins were identified, aligned, and concatenated for downstream phylogenetic inference. The phylogenetic tree was reconstructed using a maximum likelihood approach, with the best-fit substitution model selected for each gene partition in IQ-TREE v.1.6.12. The final analysis was based on 604,857 aligned sites, with 588,555 distinct patterns contributing to the tree topology.

### Methylation and RNA-seq analyses

PacBio HiFi reads with kinetic data were processed using Jasmine v3.0.0 ^117^. The output, which identified 5mC sites, was saved as ML and MM tags in the unaligned BAM file. The reads were then aligned to the *G. margarita* genome using pbmm2 1.10.0 ^118^ with the -preset HiFi parameter. PacBio CpG tools ^119^ was then used to calculate CpG methylation frequency, using the options pileup_calling_model. v1, tflite model, and –min-mapq 0. The resulting CpG locations were then intersected with genome-wide compartments, genes, and repeat regions using bedtools v2.30.0 ^120^ to determine median methylation frequencies within 50-kb windows.

For gene expression analysis, RNA-seq datasets from *G. margarita* germinating spores, ERM, and intra-radical mycelium colonizing *L. japonicus* roots were mapped to the transcriptome using Salmon v.1.3.0 ^121^. The transcriptome was first indexed using the salmon index module with the - keepDuplicates option, and reads were quantified with salmon quant, specifying the - validateMappings parameter. The Transcripts Per Million values from the salmon output were log-transformed and compared between A/B compartments across the three RNA-seq datasets.

### Compartment and topologically associating domains (TADs) analysis

HiC-Pro v2.11.1 ^122^ and HicExplorer were used to process, analyze, and visualize Hi-C data. First, HiC-Pro was used to map and filter +*Ca*Gg and *-Ca*Gg Hi-C reads, retaining only alignments with a MAPQ score greater than 10. Contact matrices were then generated at multiple resolutions, ranging from 20 kb to 100 kb in 10 kb increments. The resulting contact matrices were converted to HDF5 (.h5) format using the hicConvertFormat module in HicExplorer. The eigenvector decomposition implemented in the hicPCA command of Hicexplorer was used to call A/B compartments from the contact maps. The first eigenvector (PC1) corresponded to the A and B compartments, and the direction of the PC1 values (positive or negative) was used to determine the compartment identity. Specifically, the contact matrices were manually examined per chromosomal scaffold, and regions with identical positive values were labelled “1”, while those with negative values were labelled “2”. Finally, to assign A/B compartments, the label associated with higher average RNA-seq gene expression and gene density was assigned to compartment A, while the opposite label was assigned to compartment B.

To assess whether the presence of an endobacteria primes compartment switches, eigenvalues from contact matrices of *G. margarita* spores with and without endobacteria were compared. Regions that showed a change in PC1 values from positive to negative, or vice versa, were considered compartment switches. Additionally, the correlation of eigenvalues between the two conditions was calculated to determine the degree of shifts in compartmentalization.

We predicted TAD bodies and boundaries using hicFindTADs from HiCExplorer at 30 kb resolution, with the following parameters: --correctForMultipleTesting bonferroni and --thresholdComparisons 0.01. We then assessed the distribution of repetitive elements, protein-coding genes, and methylation levels around TAD boundaries and within ±50 kb from these boundaries using BEDTools. The resulting metaplots were generated using deepTools ^123^. Next, we compared gene expression (TPMs) patterns of boundary-associated genes and non-boundary genes across three conditions: germinating spores, ERM, and *in planta* (lotus). Additionally, to explore whether genes within the same TAD show coordinated expression, we calculated the coefficient of variation for TAD and non-TAD regions.

### *Ca*Gg genomic analyses

To confirm the number and circularity of the *Ca*Gg contigs, we reassembled HiFi reads using Flye v2.9.6^63^ and Unicycler v0.5.1^64^ genome assemblers. This process resulted in three *Ca*Gg contigs, all of which were verified as circular. One contig, containing core bacterial genes, was identified as the chromosome, while two smaller contigs containing plasmid-associated genes were classified as plasmids. The *Ca*Gg circular genome map was generated using GenoVi v0.2.16^124^. To identify Type IV Secretion System (T4SS) and Mobilization enzymes (MOB), SecReT4.0 (https://bioinfo-mml.sjtu.edu.cn/SecReT4/) and MOBscan (https://castillo.dicom.unican.es/mobscan/) were employed. Prophage regions were located using Phaster (https://phaster.ca/) with default settings. Phaster categorizes predicted prophage sequences as “intact” (score ≥ 90), “Questionable” (score 70-90), and “Incomplete” (score < 70), based on size, the presence of phage-related genes, and similarity to known phages. Predicted protein sequences were assigned KEGG Orthologs (KOs) and mapped to KEGG pathways with KofamKOALA (www.genome.jp/tools/kofamkoala/). The type II TA system was predicted using TAfinder 2.0 (https://bioinfo-mml.sjtu.edu.cn/TADB3/TAfinder.php)^125^. CRISPRCasFinder v4.2.20, implemented in Prokesee, was used to identify CRISPR arrays and cas genes.

## Acknowledgements

Our research is funded by the Discovery Program of the Natural Sciences and Engineering Research Council (RGPIN-2020-05643) and by a Discovery Accelerator Supplements Program (RGPAS-2020-00033). N.C. is a University of Ottawa Research Chair in Microbial Genomics. J.O. and G.K. were supported by MITACS projects (IT30302). At the University of Torino, research was funded under the National Recovery and Resilience Plan (NRRP), Mission 4, Component 2, Investment 1.4 - Call for tender No. 3138 of 16 December 2021, rectified by Decree n.3175 of 18 December 2021 of the Italian Ministry of University and Research, funded by the European Union – NextGenerationEU; Project code CN_00000033, Concession Decree No. 1034 of 17 June 2022, adopted by the Italian Ministry of University and Research, CUP D13C22001350001, Project title “National Biodiversity Future Center - NBFC”.

## Data Availability

The genome data used in our study are available in GenBank under the BioProject PRJNA1364746. Chromosome annotations are available at Zenodo: https://doi.org/10.5281/zenodo.18236849. All the scripts used can be accessed here: https://github.com/kenmurithi/Mugambi-2026-AMF-endosymbiont-genomics

## References

1. Spatafora, J. W. et al. A phylum-level phylogenetic classification of zygomycete fungi based on genome-scale data. Mycologia 108, 1028–1046 (2016).

2. Smith, S. E. & Read, D. J. Mycorrhizal Symbiosis. (Academic, 2008).

3. Luginbuehl, L. H. et al. Fatty acids in arbuscular mycorrhizal fungi are synthesized by the host plant. Science 356, 1175–1178 (2017).

4. Keymer, A. et al. Lipid transfer from plants to arbuscular mycorrhiza fungi. Elife 6, (2017).

5. Bonfante, P. The future has roots in the past: the ideas and scientists that shaped mycorrhizal research. New Phytol. 220, 982–995 (2018).

6. Terry, V. et al. Mycorrhizal response of Solanum tuberosum to homokaryotic versus dikaryotic arbuscular mycorrhizal fungi. Mycorrhiza (2023) doi:10.1007/s00572-023-01123-7.

7. MacColl, K. A. & Maherali, H. The effect of ecological restoration on mutualistic services provided by arbuscular mycorrhizal fungi depends on site location and host identity. Plant Soil 512, 347–360 (2025).

8. Ferguson, R. et al. Arbuscular mycorrhizal fungal genotype and nuclear organization as driving factors in host plant nutrient acquisition and stable carbon storage. Plants People Planet (2025) doi:10.1002/ppp3.10645.

9. Pozo, M. J. & Azcón-Aguilar, C. Unraveling mycorrhiza-induced resistance. Curr. Opin. Plant Biol. 10, 393–398 (2007).

10. Kokkoris, V., Stefani, F., Dalpé, Y., Dettman, J. & Corradi, N. Nuclear dynamics in the arbuscular mycorrhizal fungi. Trends Plant Sci. 25, 765–778 (2020).

11. Halary, S. et al. Conserved meiotic machinery in Glomus spp., a putatively ancient asexual fungal lineage. Genome Biol. Evol. 3, 950–958 (2011).

12. Tisserant, E. et al. Genome of an arbuscular mycorrhizal fungus provides insight into the oldest plant symbiosis. PNAS 110, 20117–20122 (2013).

13. Riley, R. & Corradi, N. Searching for clues of sexual reproduction in the genomes of arbuscular mycorrhizal fungi. Fungal Ecol. 6, 44–49 (2013).

14. Halary, S. et al. Mating Type Gene Homologues and Putative Sex Pheromone-Sensing Pathway in Arbuscular Mycorrhizal Fungi, a Presumably Asexual Plant Root Symbiont. PLoS One 8, e80729 (2013).

15. Riley, R. et al. Extreme diversification of the mating type-high-mobility group (MATA-HMG) gene family in a plant-associated arbuscular mycorrhizal fungus. New Phytol. 201, 254–268 (2014).

16. Morin, E. et al. Comparative genomics of *Rhizophagus irregularis*, *R. cerebriforme*, *R. diaphanus* and *Gigaspora rosea* highlights specific genetic features in Glomeromycotina. New Phytol. 222, 1584–1598 (2019).

17. Ropars, J. et al. Evidence for the sexual origin of heterokaryosis in arbuscular mycorrhizal fungi. Nat. Microbiol. 1, 16033 (2016).

18. Sperschneider, J. et al. Arbuscular mycorrhizal fungi heterokaryons have two nuclear populations with distinct roles in host-plant interactions. Nat. Microbiol. 8, 2142–2153 (2023).

19. Wallen, R. M. & Perlin, M. H. An overview of the function and maintenance of sexual reproduction in dikaryotic fungi. Front. Microbiol. 9, 503 (2018).

20. Ropars, J. et al. Sex in cheese: evidence for sexuality in the fungus *Penicillium roqueforti*. PLoS One 7, e49665 (2012).

21. Oliveira, J., Yildirir, G. & Corradi, N. From chaos comes order: Genetics and genome biology of arbuscular mycorrhizal fungi. Annu. Rev. Microbiol. 78, 147–168 (2024).

22. Sahraei, S. E. et al. Whole genome analyses based on single, field collected spores of the arbuscular mycorrhizal fungus *Funneliformis geosporum*. Mycorrhiza 32, 361–371 (2022).

23. Chen, E. C. H. et al. High intraspecific genome diversity in the model arbuscular mycorrhizal symbiont *Rhizophagus irregularis*. New Phytol. 220, 1161–1171 (2018).

24. Yildirir, G. et al. Long reads and Hi-C sequencing illuminate the two-compartment genome of the model arbuscular mycorrhizal symbiont *Rhizophagus irregularis*. New Phytol. 233, 1097–1107 (2022).

25. Oliveira, J. I. N. et al. Analyses of transposable elements in arbuscular mycorrhizal fungi support evolutionary parallels with filamentous plant pathogens. Genome Biol. Evol. 17, evaf038 (2025).

26. Kloppholz, S., Kuhn, H. & Requena, N. A secreted fungal effector of *Glomus intraradices* promotes symbiotic biotrophy. Curr. Biol. 21, 1204–1209 (2011).

27. Oliveira, J. I. N. & Corradi, N. Strain-specific evolution and host-specific regulation of transposable elements in the model plant symbiont *Rhizophagus irregularis*. G3 (Bethesda) 14, jkae055 (2024).

28. Teulet, A. et al. A pathogen effector FOLD diversified in symbiotic fungi. New Phytol. 239, 1127–1139 (2023).

29. Lanfranco, L. & Bonfante, P. Lessons from arbuscular mycorrhizal fungal genomes. Curr. Opin. Microbiol. 75, 102357 (2023).

30. Torres, D. E., Reckard, A. T., Klocko, A. D. & Seidl, M. F. Nuclear genome organization in fungi: from gene folding to Rabl chromosomes. FEMS Microbiol. Rev. 47, fuad021 (2023).

31. Dixon, J. R. et al. Topological domains in mammalian genomes identified by analysis of chromatin interactions. Nature 485, 376–380 (2012).

32. Dekker, J. & Heard, E. Structural and functional diversity of Topologically Associating Domains. FEBS Lett. 589, 2877–2884 (2015).

33. Glavincheska, I. & Lorrain, C. Three-dimensional genome architecture connects chromatin structure and function in a major wheat pathogen. *<u>bioRxiv</u>* 2025.05.13.653796 (2025) doi:10.1101/2025.05.13.653796.

34. Kurbidaeva, A. et al. Topologically associating domains and the evolution of three-dimensional genome architecture in rice. Plant J. 122, e70139 (2025).

35. Winter, D. J. et al. Repeat elements organise 3D genome structure and mediate transcription in the filamentous fungus *Epichloë festucae*. PLoS Genet. 14, e1007467 (2018).

36. Zhang, G., Li, Y. & Wei, G. Multi-omic analysis reveals dynamic changes of three-dimensional chromatin architecture during T cell differentiation. *Commun*. Biol. 6, 773 (2023).

37. Hansen, A. S., Cattoglio, C., Darzacq, X. & Tjian, R. Recent evidence that TADs and chromatin loops are dynamic structures. Nucleus 9, 20–32 (2018).

38. Li, H., Playter, C., Das, P. & McCord, R. P. Chromosome compartmentalization: causes, changes, consequences, and conundrums. Trends Cell Biol. 34, 707–727 (2024).

39. Venice, F. et al. At the nexus of three kingdoms: the genome of the mycorrhizal fungus *Gigaspora margarita* provides insights into plant, endobacterial and fungal interactions. Environ. Microbiol. 22, 122–141 (2020).

40. Bonfante, P. & Desirò, A. Who lives in a fungus? The diversity, origins and functions of fungal endobacteria living in Mucoromycota. ISME J. 11, 1727–1735 (2017).

41. Turina, M. et al. The virome of the arbuscular mycorrhizal fungus *Gigaspora margarita* reveals the first report of DNA fragments corresponding to replicating non-retroviral RNA viruses in fungi. Environ. Microbiol. 20, 2012–2025 (2018).

42. Salvioli, A. et al. Symbiosis with an endobacterium increases the fitness of a mycorrhizal fungus, raising its bioenergetic potential. ISME J. 10, 130–144 (2016).

43. Cheng, H., Concepcion, G. T., Feng, X., Zhang, H. & Li, H. Haplotype-resolved de novo assembly using phased assembly graphs with hifiasm. Nat. Methods 18, 170–175 (2021).

44. Tang, N. et al. A survey of the gene repertoire of *Gigaspora rosea* unravels conserved features among Glomeromycota for obligate biotrophy. Front. Microbiol. 7, 233 (2016).

45. Rosling, A. et al. Evolutionary history of arbuscular mycorrhizal fungi and genomic signatures of obligate symbiosis. BMC Genomics 25, 529 (2024).

46. Ghignone, S. et al. The genome of the obligate endobacterium of an AM fungus reveals an interphylum network of nutritional interactions. ISME J. 6, 136–145 (2012).

47. Malar C, M. et al. Early branching arbuscular mycorrhizal fungus *Paraglomus occultum* carries a small and repeat-poor genome compared to relatives in the Glomeromycotina. Microb. Genom. 8, 000810 (2022).

48. Malar C, M. et al. The genome of *Geosiphon pyriformis* reveals ancestral traits linked to the emergence of the arbuscular mycorrhizal symbiosis. Curr. Biol. 31, 1578–1580 (2021).

49. Pelin, A. et al. The mitochondrial genome of the arbuscular mycorrhizal fungus *Gigaspora margarita* reveals two unsuspected trans-splicing events of group I introns. New Phytol. 194, 836–845 (2012).

50. Wijayawardene, N. N. et al. Outline of fungi and fungus-like taxa – 2021. Mycosphere 13, 53–453 (2022).

51. Beaudet, D. et al. Ultra-low input transcriptomics reveal the spore functional content and phylogenetic affiliations of poorly studied arbuscular mycorrhizal fungi. DNA Res. 0, 1–11 (2017).

52. Błaszkowski, J. et al. A new order, Entrophosporales, and three new Entrophospora species in Glomeromycota. Front. Microbiol. 13, 962856 (2022).

53. Redecker, D. et al. An evidence-based consensus for the classification of arbuscular mycorrhizal fungi (Glomeromycota). Mycorrhiza (2013) doi:10.1007/s00572-013-0486-y.

54. Corradi, N., Antunes, P. M. & Magurno, F. A call for reform: implementing genome-based approaches for species classification in Glomeromycotina. New Phytol. (2025) doi:10.1111/nph.70148.

55. Maeda, T. et al. Evidence of non-tandemly repeated rDNAs and their intragenomic heterogeneity in *Rhizophagus irregularis*. *Commun*. Biol. 1, 87 (2018).

56. Stefani, F. et al. The pitfalls of rDNA-based AMF identification: a comparative analysis of rDNA and protein-coding genes. New Phytol. 248, 1501–1515 (2025).

57. Boisvert, F.-M., van Koningsbruggen, S., Navascués, J. & Lamond, A. I. The multifunctional nucleolus. Nat. Rev. Mol. Cell Biol. 8, 574–585 (2007).

58. Pederson, T. The nucleolus. Cold Spring Harb. Perspect. Biol. 3, a000638–a000638 (2011).

59. Schöfer, C. & Weipoltshammer, K. Nucleolus and chromatin. Histochem. Cell Biol. 150, 209–225 (2018).

60. Muszewska, A., Steczkiewicz, K., Stepniewska-Dziubinska, M. & Ginalski, K. Transposable elements contribute to fungal genes and impact fungal lifestyle. Sci. Rep. (2019) doi:10.1038/s41598-019-40965-0.

61. Dallaire, A. et al. Transcriptional activity and epigenetic regulation of transposable elements in the symbiotic fungus *Rhizophagus irregularis*. Genome Res. 31, 2290–2302 (2021).

62. Li, D. et al. Comparative 3D genome architecture in vertebrates. BMC Biol. 20, 99 (2022).

63. Kolmogorov, M., Yuan, J., Lin, Y. & Pevzner, P. A. Assembly of long, error-prone reads using repeat graphs. Nat. Biotechnol. 37, 540–546 (2019).

64. Wick, R. R., Judd, L. M., Gorrie, C. L. & Holt, K. E. Unicycler: Resolving bacterial genome assemblies from short and long sequencing reads. PLoS Comput. Biol. 13, e1005595 (2017).

65. Sorwar, E., Oliveira, J. I. N., Malar C, M., Krüger, M. & Corradi, N. Assembly and comparative analyses of the *Geosiphon pyriformis* metagenome. Environ. Microbiol. 26, e16681 (2024).

66. Chan, W. T., Garcillán-Barcia, M. P., Yeo, C. C. & Espinosa, M. Type II bacterial toxin-antitoxins: hypotheses, facts, and the newfound plethora of the PezAT system. FEMS Microbiol. Rev. 47, fuad052 (2023).

67. Salvioli di Fossalunga, A., Lipuma, J., Venice, F., Dupont, L. & Bonfante, P. The endobacterium of an arbuscular mycorrhizal fungus modulates the expression of its toxin-antitoxin systems during the life cycle of its host. ISME J. 11, 2394–2398 (2017).

68. Teulet, A., et al. A pathogen effector FOLD diversified in symbiotic fungi. *bioRxiv* (2022) doi:10.1101/2022.12.16.520752.

69. Voß, S., Betz, R., Heidt, S., Corradi, N. & Requena, N. RiCRN1, a crinkler effector from the arbuscular mycorrhizal fungus *Rhizophagus irregularis*, functions in arbuscule development. Front. Microbiol. 9, 2068 (2018).

70. Bruns, T. D., Corradi, N., Redecker, D., Taylor, J. W. & Öpik, M. Glomeromycotina: What is a species and why should we care? New Phytol. (2017) doi:10.1111/nph.14913.

71. Reinhardt, D., Roux, C., Corradi, N. & Di Pietro, A. Lineage-Specific Genes and Cryptic Sex: Parallels and Differences between Arbuscular Mycorrhizal Fungi and Fungal Pathogens. Trends Plant Sci. (2020) doi:10.1016/j.tplants.2020.09.006.

72. Dong, S., Raffaele, S. & Kamoun, S. The two-speed genomes of filamentous pathogens: waltz with plants. Curr. Opin. Genet. Dev. 35, 57–65 (2015).

73. Manley, B. F. et al. A highly contiguous genome assembly reveals sources of genomic novelty in the symbiotic fungus *Rhizophagus irregularis*. G3 (Bethesda) 13, jkad077 (2023).

74. Dixon, J. R., Gorkin, D. U. & Ren, B. Chromatin domains: The unit of chromosome organization. Mol. Cell 62, 668–680 (2016).

75. Grummt, I. Life on a planet of its own: regulation of RNA polymerase I transcription in the nucleolus. Genes Dev. 17, 1691–1702 (2003).

76. Mancio-Silva, L., Zhang, Q., Scheidig-Benatar, C. & Scherf, A. Clustering of dispersed ribosomal DNA and its role in gene regulation and chromosome-end associations in malaria parasites. Proc. Natl. Acad. Sci. U. S. A. 107, 15117–15122 (2010).

77. Rabuffo, C. et al. Inter-chromosomal transcription hubs shape the 3D genome architecture of African trypanosomes. Nat. Commun. 15, 10716 (2024).

78. Jargeat, P. et al. Isolation, free-living capacities, and genome structure of “*Candidatus* Glomeribacter gigasporarum,” the endocellular bacterium of the mycorrhizal fungus *Gigaspora margarita*. J. Bacteriol. 186, 6876–6884 (2004).

79. Pawlowska, T. E. et al. Biology of fungi and their bacterial endosymbionts. Annu. Rev. Phytopathol. 56, 289–309 (2018).

80. Uehling, J. K. et al. Bacterial endosymbionts of Mucoromycota fungi: Diversity and function of their interactions. in The Mycota 177–205 (Springer International Publishing, Cham, 2023).

81. Winsor, G. L. et al. The Burkholderia Genome Database: facilitating flexible queries and comparative analyses. Bioinformatics 24, 2803–2804 (2008).

82. Strübing, U., Lucius, R., Hoerauf, A. & Pfarr, K. M. Mitochondrial genes for heme-dependent respiratory chain complexes are up-regulated after depletion of Wolbachia from filarial nematodes. Int. J. Parasitol. 40, 1193–1202 (2010).

83. Venice, F., et al. *Gigaspora margarita* with and without its endobacterium shows adaptive responses to oxidative stress. Mycorrhiza 27, 747–759 (2017).

84. Wang, J. et al. Crucial involvement of heme biosynthesis in vegetative growth, development, stress response, and fungicide sensitivity of *Fusarium graminearum*. Int. J. Mol. Sci. 25, 5268 (2024).

85. Horvath, P. & Barrangou, R. CRISPR/Cas, the immune system of bacteria and archaea. Science 327, 167–170 (2010).

86. Rostøl, J. T. & Marraffini, L. (ph)ighting phages: How bacteria resist their parasites. Cell Host Microbe 25, 184–194 (2019).

87. Burstein, D. et al. Major bacterial lineages are essentially devoid of CRISPR-Cas viral defence systems. Nat. Commun. 7, 10613 (2016).

88. Siozios, S. et al. Genome dynamics across the evolutionary transition to endosymbiosis. Curr. Biol. 34, 5659–5670.e7 (2024).

89. Song, S. & Wood, T. K. A primary physiological role of toxin/antitoxin systems is phage inhibition. Front. Microbiol. 11, 1895 (2020).

90. Lumini, E. et al. Presymbiotic growth and sporal morphology are affected in the arbuscular mycorrhizal fungus *Gigaspora margarita* cured of its endobacteria. Cell. Microbiol. 9, 1716–1729 (2007).

91. Spores of mycorrhizal Endogone species extracted from soil by wet sieving and decanting. Transactions of the British Mycological Society 46, 235–244 (1963).

92. Wolff, J. et al. Galaxy HiCExplorer 3: a web server for reproducible Hi-C, capture Hi-C and single-cell Hi-C data analysis, quality control and visualization. Nucleic Acids Res. 48, W177–W184 (2020).

93. Cheng, H. et al. Haplotype-resolved assembly of diploid genomes without parental data. Nat. Biotechnol. 40, 1332–1335 (2022).

94. Uliano-Silva, M. et al. MitoHiFi: a python pipeline for mitochondrial genome assembly from PacBio high fidelity reads. BMC Bioinformatics 24, 288 (2023).

95. Schwengers, O. et al. Bakta: rapid and standardized annotation of bacterial genomes via alignment-free sequence identification. Microb. Genom. 7, (2021).

96. Arima Genomics. Mapping_pipeline. (2019).

97. Zhou, C., McCarthy, S. A. & Durbin, R. YaHS: yet another Hi-C scaffolding tool. Bioinformatics 39, (2023).

98. Harry, E. PretextView (Paired REad TEXTure Viewer): A Desktop Application for Viewing Pretext Contact Maps. (Github, 2020).

99. Durand, N. C. et al. Juicer provides a one-click system for analyzing loop-resolution Hi-C experiments. Cell Syst. 3, 95–98 (2016).

100. Hao, Z. et al. RIdeogram: drawing SVG graphics to visualize and map genome-wide data on the idiograms. PeerJ Comput. Sci. 6, e251 (2020).

101. Flynn, J. M. et al. RepeatModeler2 for automated genomic discovery of transposable element families. Proc. Natl. Acad. Sci. U. S. A. 117, 9451–9457 (2020).

102. Qian, J. et al. TEtrimmer: a tool to automate the manual curation of transposable elements. Nat. Commun. 16, 8429 (2025).

103. Smit, A. F. A., Hubley, R. & Green, P. RepeatMasker. *Published on the web at* http://www.repeatmasker.org (1996).

104. Venice, F. et al. Symbiotic responses of *Lotus japonicus* to two isogenic lines of a mycorrhizal fungus differing in the presence/absence of an endobacterium. Plant J. 108, 1547–1564 (2021).

105. Jin, Y., Tam, O. H., Paniagua, E. & Hammell, M. TEtranscripts: a package for including transposable elements in differential expression analysis of RNA-seq datasets. Bioinformatics 31, 3593–3599 (2015).

106. Palmer, J. M. & Stajich, J. Funannotate v1.8.1: Eukaryotic Genome Annotation. (Zenodo, 2020). doi:10.5281/ZENODO.4054262.

107. Simão, F. A., Waterhouse, R. M., Ioannidis, P., Kriventseva, E. V. & Zdobnov, E. M. BUSCO: assessing genome assembly and annotation completeness with single-copy orthologs. Bioinformatics 31, 3210–3212 (2015).

108. Teufel, F. et al. SignalP 6.0 predicts all five types of signal peptides using protein language models. Nat. Biotechnol. 40, 1023–1025 (2022).

109. Sperschneider, J. & Dodds, P. N. EffectorP 3.0: Prediction of apoplastic and cytoplasmic effectors in fungi and Oomycetes. Mol. Plant. Microbe. Interact. 35, 146–156 (2022).

110. Huang, L. et al. dbCAN-seq: a database of carbohydrate-active enzyme (CAZyme) sequence and annotation. Nucleic Acids Res. 46, D516–D521 (2018).

111. Zheng, J. et al. dbCAN3: automated carbohydrate-active enzyme and substrate annotation. Nucleic Acids Res. 51, W115–W121 (2023).

112. Orłowska, M., Steczkiewicz, K. & Muszewska, A. Utilization of cobalamin is ubiquitous in early-branching fungal phyla. Genome Biol. Evol. 13, evab043 (2021).

113. Katoh, K. & Standley, D. M. MAFFT multiple sequence alignment software version 7: improvements in performance and usability. Mol. Biol. Evol. 30, 772–780 (2013).

114. Miller, M. A., Pfeiffer, W. & Schwartz, T. The CIPRES science gateway: a community resource for phylogenetic analyses. in *Proceedings of the 2011 TeraGrid Conference: Extreme Digital Discovery* (ACM, New York, NY, USA, 2011). doi:10.1145/2016741.2016785.

115. Minh, B. Q. et al. IQ-TREE 2: New models and efficient methods for phylogenetic inference in the genomic era. Mol. Biol. Evol. 37, 1530–1534 (2020).

116. Magurno, F., et al. *Glomus mongioiense*, a new species of arbuscular mycorrhizal fungi from Italian Alps and the phylogeny-spoiling issue of ribosomal variants in the *Glomus* genus. Agronomy (Basel) 14, 1350 (2024).

117. jasmine. Jasmine: Call Select Base Modifications in PacBio HiFi Reads. (Github, 2023).

118. pbmm2. Pbmm2: A Minimap2 Frontend for PacBio Native Data Formats. (Github, 2017).

119. pb-CpG-tools. Pb-CpG-Tools: Collection of Tools for the Analysis of CpG Data. (Github, 2022).

120. Quinlan, A. R. & Hall, I. M. BEDTools: a flexible suite of utilities for comparing genomic features. Bioinformatics 26, 841–842 (2010).

121. Patro, R., Duggal, G., Love, M. I., Irizarry, R. A. & Kingsford, C. Salmon provides fast and bias-aware quantification of transcript expression. Nat. Methods 14, 417–419 (2017).

122. Servant, N. et al. HiC-Pro: an optimized and flexible pipeline for Hi-C data processing. Genome Biol. 16, 259 (2015).

123. Ramírez, F., Dündar, F., Diehl, S., Grüning, B. A. & Manke, T. deepTools: a flexible platform for exploring deep-sequencing data. Nucleic Acids Res 42, W187–W191 (2014).

124. Cumsille, A. et al. GenoVi, an open-source automated circular genome visualizer for bacteria and archaea. PLoS Comput. Biol. 19, e1010998 (2023).

125. Guan, J. et al. TADB 3.0: an updated database of bacterial toxin-antitoxin loci and associated mobile genetic elements. Nucleic Acids Res. 52, D784–D790 (2024).

