## Supplemental figures for "The 3D Genome of *Gigaspora margarita* Unveils Stable Chromatin and Nucleolar Organization and Symbiont-Dependent Genome Dynamics"

**Figure S1.** Maximum likelihood phylogenetic tree showing the placement of *Gigaspora margarita* on the fungal tree of life based on 702 single-copy orthologs. Bootstrap support values are indicated on each branch.

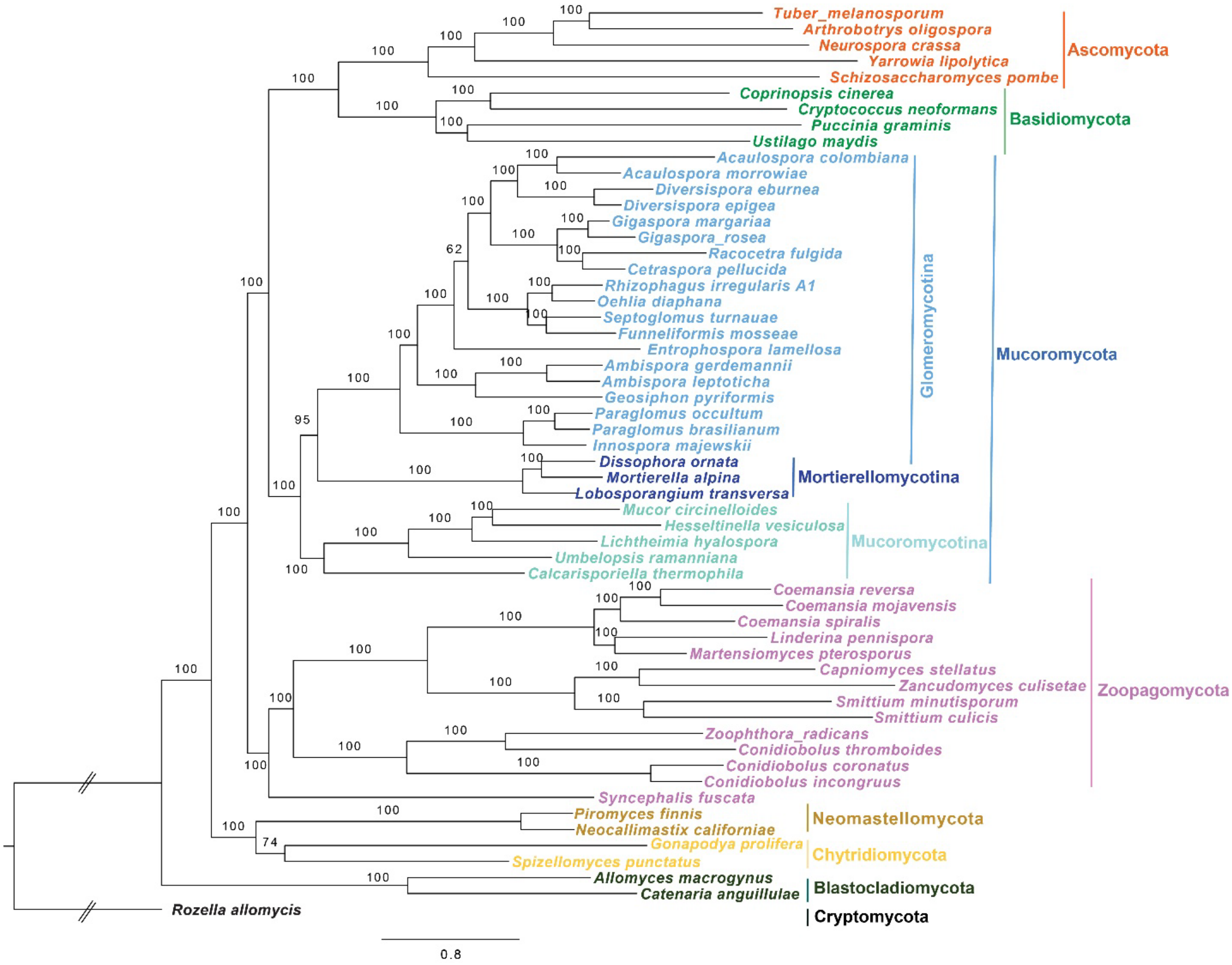

**Figure S2. Phylogram generated from Maximum Likelihood (ML) analyses showing the phylogenetic relationships among *G. margarita* and other *Gigaspora* species possessing rDNA sequences**. Sequences of *Dentiscutata savannicola* and *Fuscutata heterogama* were used to root the tree. Different colours for the external branches indicate the species shown in the legend. Internal branches are coloured to match the same species notation as their descendant leaves when those leaves are of a single species. Bootstrap values, ultrafast bootstrap values, and posterior probabilities ≥ 90, 90, and 0.98, respectively, are indicated above or below the branches. The bar represents 0.02 expected changes per site per branch.

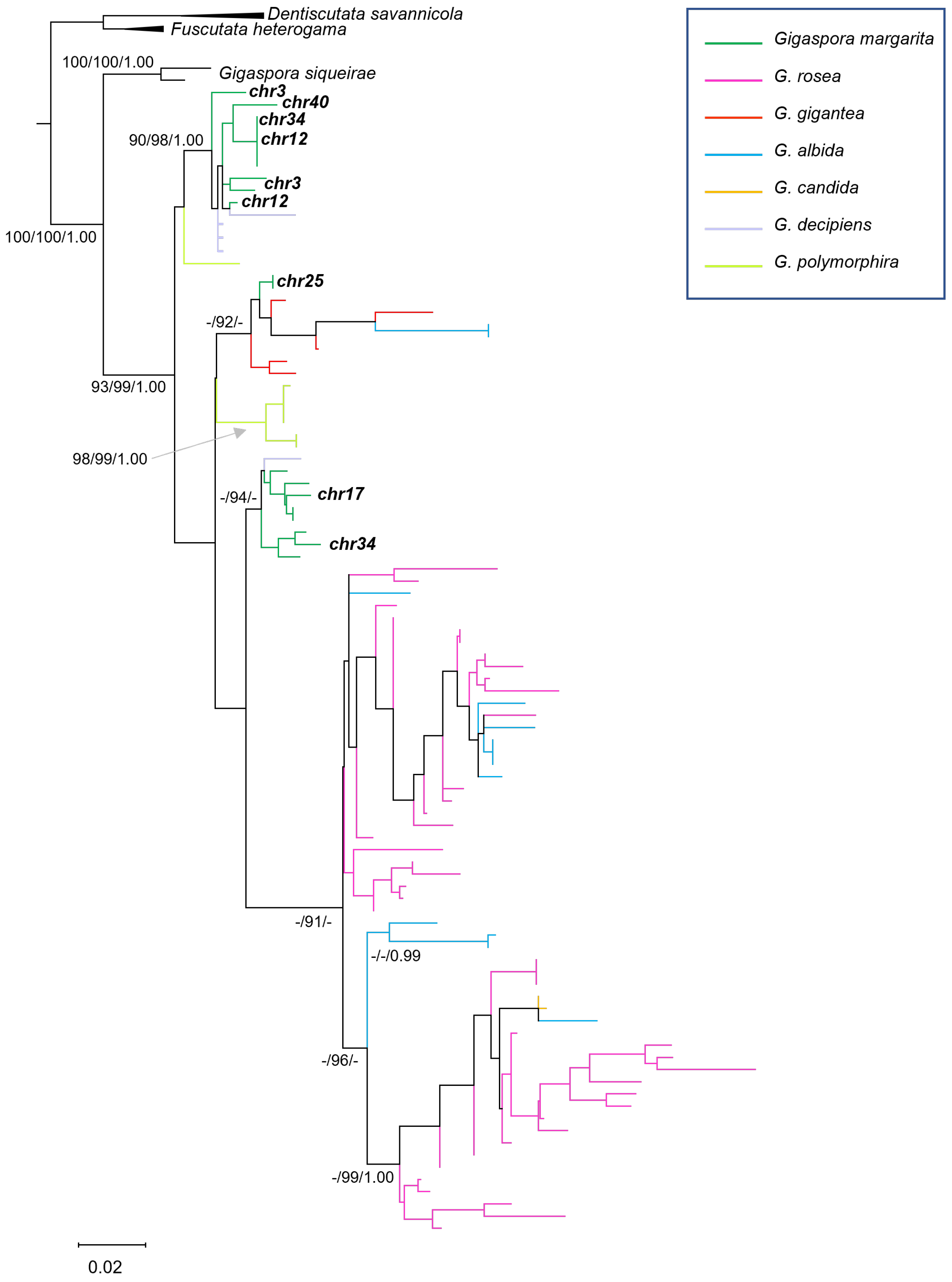

**Figure S3.** Genome-wide Hi-C contact maps of (A) *G. margarita* and (B) *R. irregularis*. The black squares represent chromosomes sorted by size. The colour intensity corresponds to interaction frequencies between loci, with darker red indicating high contact probability and white signifying low or no interactions. The circles highlight physical interactions between rDNA across different chromosomes.

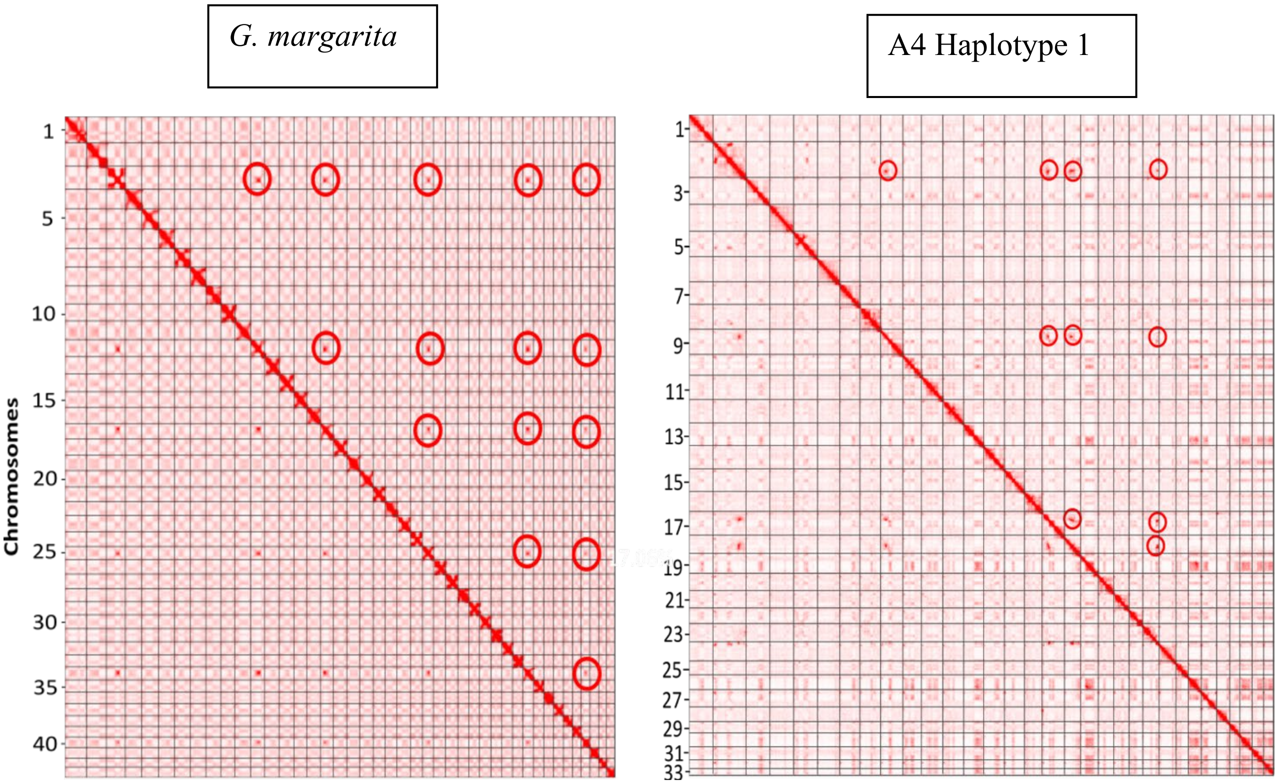

**Figure S4.** Hi-C contact maps showing A/B compartments in *G. margarita* chromosomes at a 50 kb resolution in B- and B+ conditions. Regions that interact more frequently are visualized as brighter squares on the contact map.

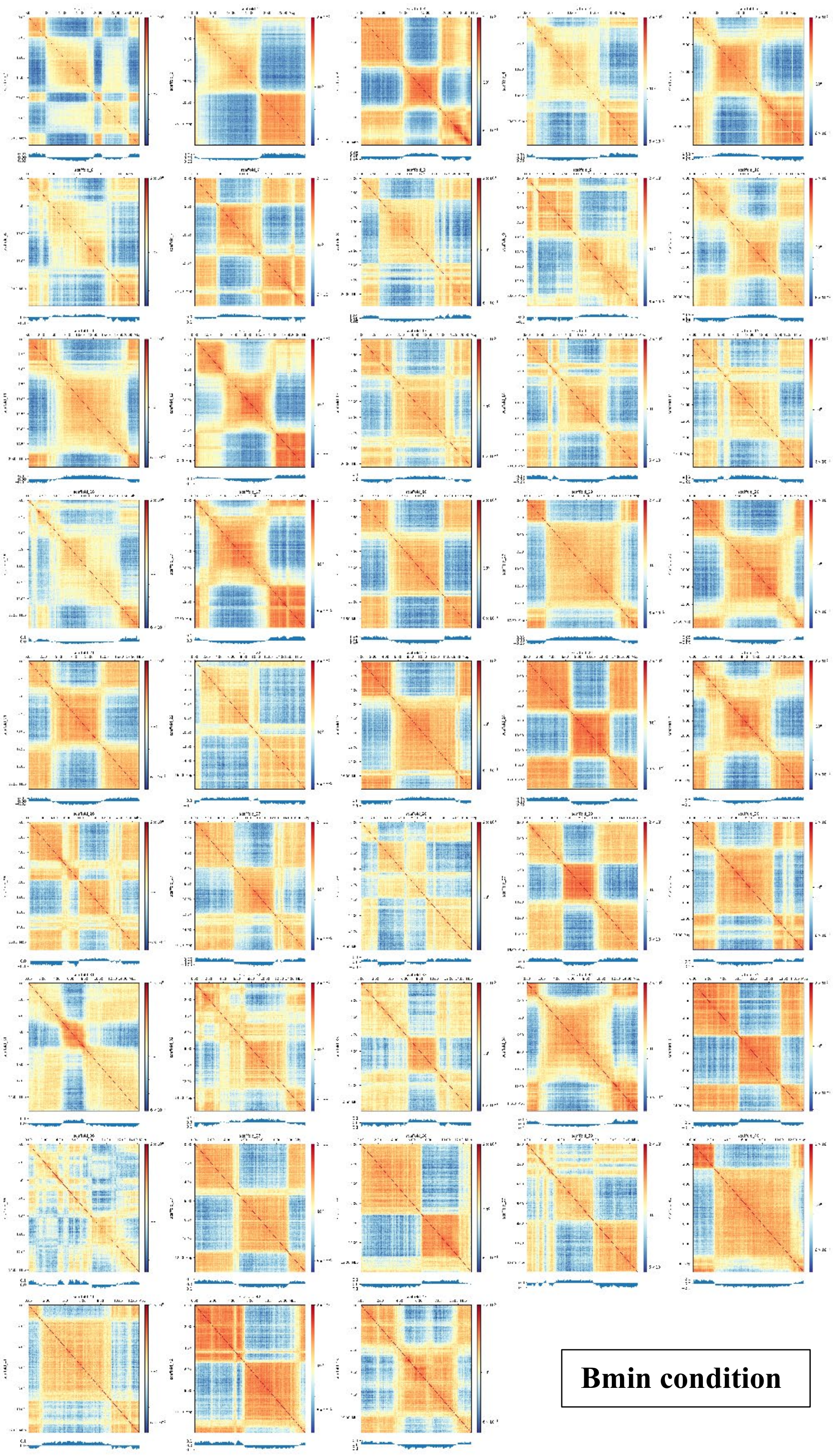

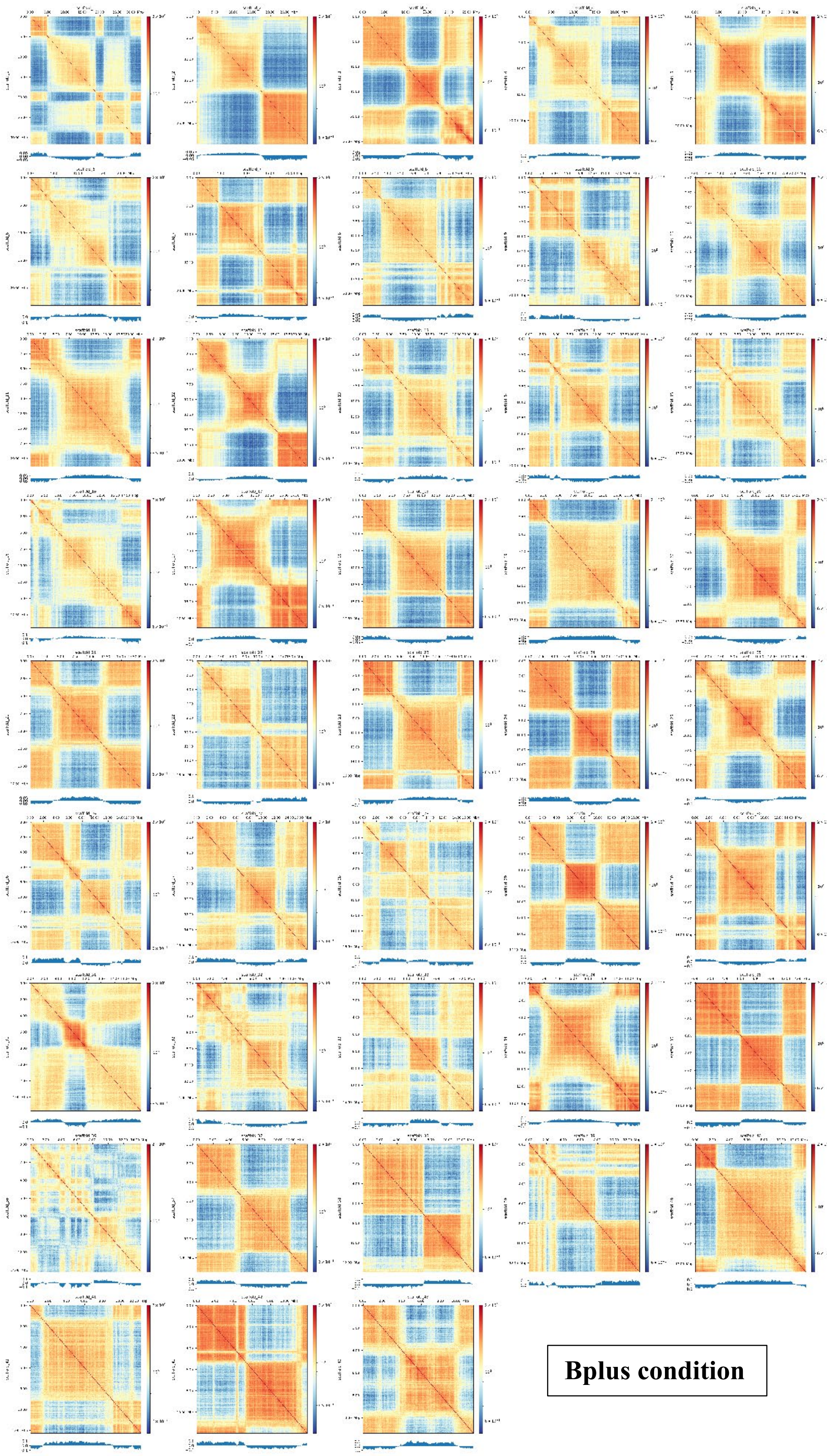

**Bplus condition**

**Figure S5.** Boxplots showing variation of expression across conditions: ERM, germinating spores, and *in planta* (lotus) within TAD and non-TAD regions. Boxes show the first quartile (25%), the middle black line (50%), and the third quartile (75%). The whiskers extend to 1.5× the box length, and the outliers are represented as dots. Asterisks above the boxplots indicate significant differences between the A/B compartments ($p < 0.05$, Wilcoxon rank-sum test).

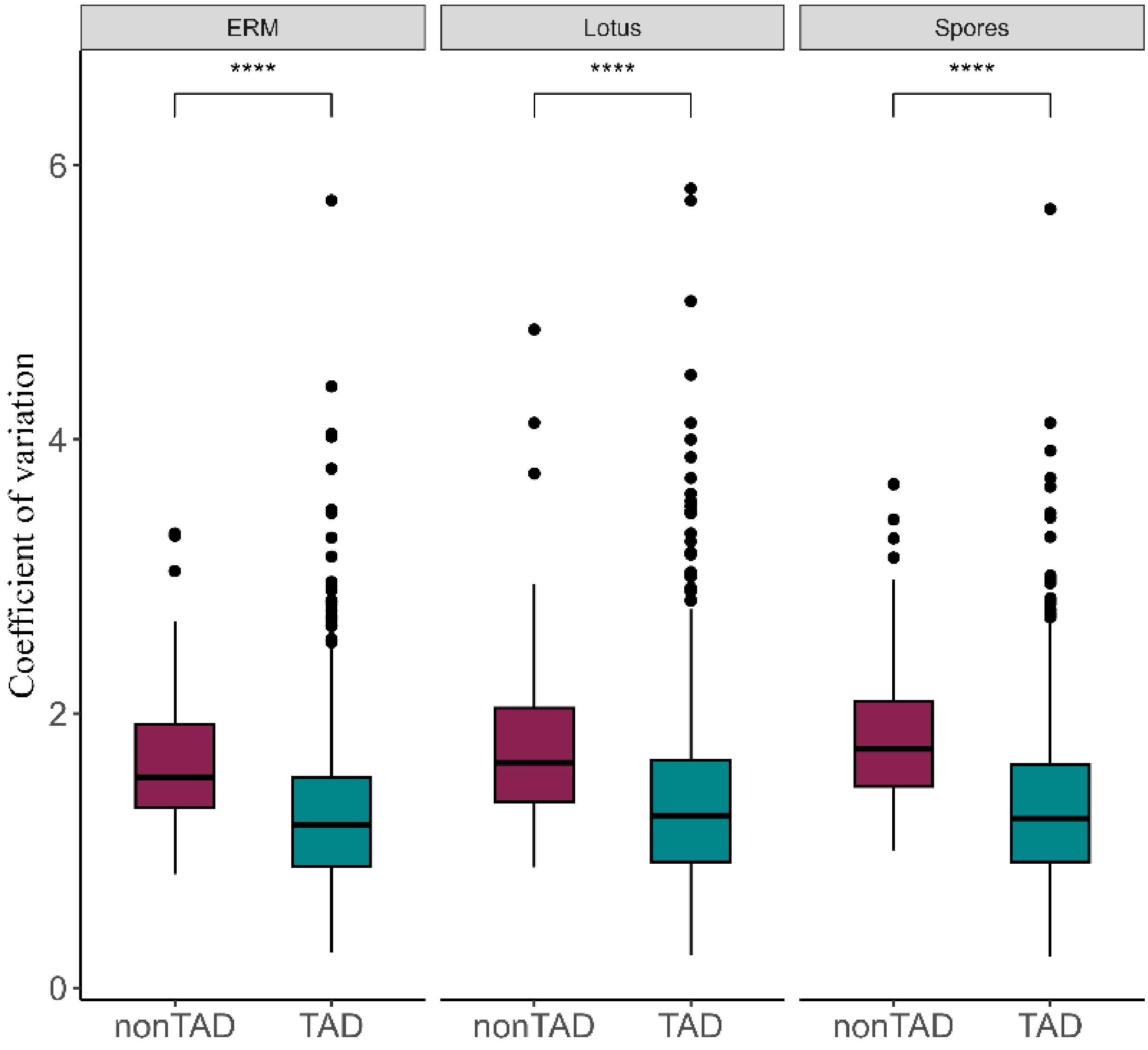

**Figure S6.** Hi-C contact maps of *Candidatus* Glomerobacter gigasporarum (*Ca*Gg) chromosome. Regions that interact more frequently are visualized as brighter squares on the contact map

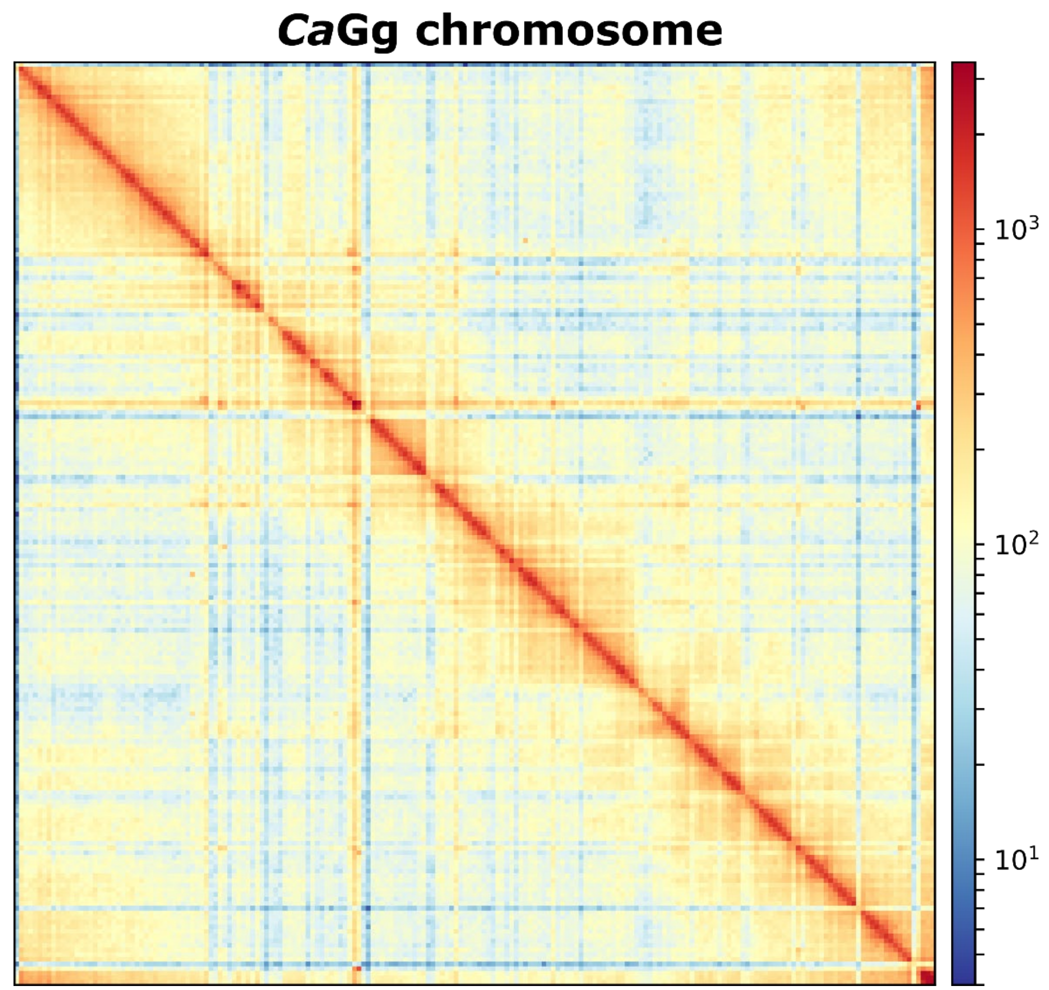
